# Exploring the role of multivariate associations between brain networks and gut microbiota in (food) value-based decision-making

**DOI:** 10.64898/2026.10.12.758555

**Authors:** Silvia Papalini, Esther Aarts, Nils Kohn

## Abstract

Food reward valuation and decision making are key psychological processes regulating eating behaviour. Stress and negative affect can shape these processes, increasing vulnerability to overeating high-caloric, ultra-processed foods and thereby elevating obesity risk. Yet not all individuals under stress exhibit such behavioural shifts, highlighting the need to understand which modulating factors explain this inter-individual variability. Emerging evidence suggests that anxiety-/stress-induced changes in gut microbiota might alter reward sensitivity via the gut–brain axis. An integrative framework linking microbiota, neural, and affective processes is thus essential for explaining individual differences in emotional eating, yet such a framework is still lacking. To this end, supervised linked independent component analysis (Super-BigFLICA) was used to explore multivariate gut–brain latent components guided by affective and body composition indexes in healthy participants (N = 592) from the Healthy Brain Study. Next, general linear models were used to identify if any of the observed components were predictive of elevated unhealthy food valuation and choice in a food auction task. A multimodal component was predictive of both food valuation and choice. This component was characterized by co-occurring changes in several gut bacterial taxa at the genus level (e.g., UCG-010 and DTU089), specific connectivity between reward-related brain regions (e.g., nucleus accumbens, caudate), and was clearly linked to elevated anxiety sensitivity, life-event impact, and body mass index. These findings demonstrate the feasibility of supervised gut-brain integration for predicting behaviour and reveal a potential phenotype linking microbiota composition, brain function, and emotional state that should be further investigated in larger cohorts characterized by emotional eating and overweight.

## 1 Introduction

Food choices are guided by value-based decision-making processes that integrate attributes such as taste and health into a common value signal that drives behaviour [1, 2]. Generally, these processes support adaptive regulation of energy intake by appropriately weighing long-term health goals against short-term rewards. However, this regulation is fragile: the same valuation mechanisms can become biased toward immediately rewarding attributes such as palatability or emotional relief, thereby increasing the risk of overweight and obesity. This vulnerability raises a critical open question, namely, under which conditions the valuation process becomes decoupled from homeostatic needs and shifts toward unhealthy options.

Stress and anxiety are prominent candidates for driving such shifts [3, 4]. Convergent evidence indicates that stress can alter both the subjective value assigned to food and the choices individuals make [5–7]. However, these effects are strikingly heterogeneous [8]. Some studies report a reduced reinforcing value of food under stress [9], while others show a higher preference for highly palatable, energy-dense foods or a selective bias towards taste over health attributes [10–12]. At the neural level, stress has been linked to altered interactions between prefrontal control systems and valuation circuits [10], suggesting that it can bias decision-making not by uniformly amplifying or suppressing reward signals, but by changing how different value attributes are weighed. This variability challenges unitary accounts of emotional overeating and suggests that additional factors shape how stress and anxiety influence food-related decisions.

One limitation of current models is their primary focus on central nervous system mechanisms. Specifically, in the context of food reward, peripheral physiological systems (particularly those associated with the gastrointestinal tract) play a central role in signaling the state of nutrient and internal bodily conditions to the brain [13, 14]. Among these, the gut microbiota has emerged as a key modulator of brain function through the gut–brain axis [15]. The composition of the microbiota is sensitive to both stress and diet and has been linked to changes in neural activity and behaviour [16]. This raises the possibility that inter-individual differences in the gut microbiota contribute to variability in the way stress shapes food valuation and choice, either by modulating the central reward circuit or by altering the physiological response of the organism to stress (or both). Despite this, the joint contributions of brain function and gut microbiota to food-related decision-making, and their modulation by stress, remain poorly understood. Addressing this question requires approaches that go beyond single-system analyses, and instead capture distributed patterns of variation across the brain, body, and behaviour.

Recent advances in multimodal data integration techniques offer several tools to bridge this gap. In particular, FMRIB’s Linked Independent Component Analysis (FLICA) [17] and its supervised extension (SuperBigFLICA) [18] represent novel approaches developed for integrating multiple imaging modalities (including the gut microbiota [19]) into a single predictive framework. Specifically, these methods allow a model to learn latent components capturing shared variance across multiple modalities such as brain imaging (e.g., resting-state and/or task-based functional magnetic resonance imaging [fMRI], brain anatomy, etc.) and gut microbiota composition. These latent components represent potential multivariate biomarkers (phenotypes) that can be used to predict behavioural outcomes using classical inference models. The supervised extension allows for tuning of multimodal integration using information from psychological traits and/or states of interest to optimize this behavioural prediction. Hence, our aim was twofold: to provide a proof-of-concept for this integrative approach; and to explore whether latent components, extracted via a decomposition guided by stress, negative affect, and body composition variables, can predict elevated unhealthy food valuation and decision-making in a non-clinical sample, capturing subclinical tendencies.

In this preregistered study, we applied SuperBigFLICA to a large sample from the Healthy Brain Study [HBS] [20]. This non-clinical cohort shows substantial variability in Body Mass Index (BMI) and is free from confounds of chronic illness and treatment history, allowing us to test the validity of this integrative approach and explore phenotypes that remain relevant to clinical models of overeating. The specific hypothesis of this preregistered study (https://osf.io/v2dep) is that the elevated value and choice of sweet and salty snacks (unhealthy food items) are predicted by hidden multivariate information between reward-related brain networks and the composition of the gut microbiota that are ‘steered’ by stress/affective psychological variables. Briefly, to test this hypothesis, we integrated resting-state fMRI connectivity from four large-scale brain networks (the Salience Network [SN], the Default Mode Network [DMN], the Executive Control Network [ECN], and the Limbic Network [LN]) with gut microbiota composition. These modality inputs represented the microbiological and Imaging-Derived Phenotypes (called IDPs, even if they also included the gut microbiota) of our interest. Supervision of the discovery of the hidden components was guided by preregistered stress-, negative-emotion-, and other related psychological, and physiological variables (called non-Imaging-Derived Phenotypes, or nIDPs). The model search involved extensive hyperparameter tuning, dimensionality reduction, and clustering to identify robust, generalizable models. The centroids of the clusters represented the selected models. From these selected models, components with a gutbrain mixture of at least 15% were used as independent variables to test conceptual validity and to predict both the valuation and choice rates of unhealthy foods from an independent Food Auction Task (FAT) dataset [21] from the same participants. This dataset was excluded from model training.

## 2 Results

### 2.1 Descriptive of the extracted sample

**Table 1** shows the means (M) and standard deviations (SD) of the demographic, subjective (e.g., questionnaires), and physiological features (nIDPs) of the participants extracted from the first assessment of HBS. The whole sample included N = 340 Females and N = 252 Males (total N = 592), aged between 30 and 39 years (M = 33.81, SD = 2.82).

| Variable name | M | SD | Count (N) |
| --- | --- | --- | --- |
| State and Trait Anxiety Scale (STS)[22] | 33.28 | 8.27 | 583 |
| Perceived Stress Scale (PSS) [23] | 12.91 | 5.84 | 541 |
| Kilocalories per day from Food Frequency Questionnaire (FFQ) (Kilokal) [24] | 1993.87 | 606.4 | 549 |
| Body fat percentage (Fat_perc) | 23.95 | 8.28 | 510 |
| Systolic Blood Pressure (Sys_BP) | 116.52 | 11.9 | 582 |
| Diastolic Blood Pressure (Dia_BP) | 69.42 | 9.54 | 582 |
| Anxiety Sensitivity Index (ASI) [25] | 11.08 | 6.19 | 529 |
| Childhood Trauma Questionnaire (CTQ) [26] | 34.35 | 7.66 | 449 |
| Influence of Life Events (LE_influence) [27] | 44.06 | 21.02 | 534 |
| Body Mass Index (BMI) | 24.60 | 3.56 | 581 |
| Waist Circumference (Waist_Cir) | 84.07 | 10.85 | 537 |
| Self-control Scale (SCO) [28] | 40.65 | 7.56 | 526 |

### 2.2 Imaging-Derived Phenotypes (IDPs) including gut microbiota

#### Brain Networks

Four out of twenty-five networks obtained through Independent Component Analysis (ICA) were selected and used as input modality; the selection of these networks was based on their relevance in (food) valuation and decision making, food craving, and ‘habitual’ food (over)consumption, see **Fig. 1** for brain activations obtained using a threshold of *Z >* 3.0.

**Fig. 1:**
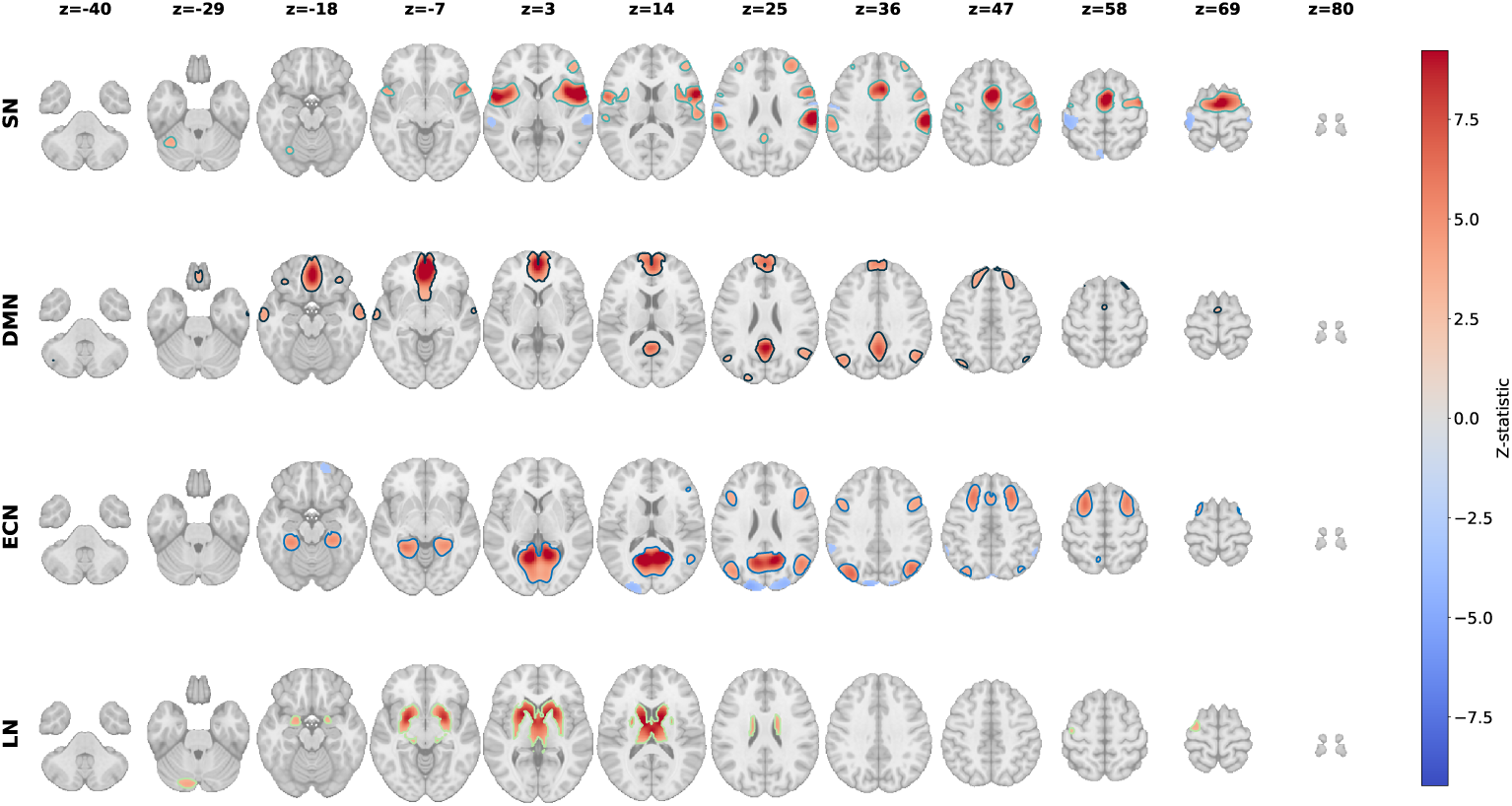
Visualization of the selected brain networks. The Salience Network [SN, with heatmap outlined in turquoise], the Default Mode Network [DMN, with heatmap outlined in dark blue], the Executive Control Network [ECN, with heatmap outlined in light blue], and the Limbic Network [LN, with heatmap outlined in light green]. Resting-state network maps were obtained using dual-regression analysis and thresholded at *Z >* 3.0. Activations are displayed on a standardized Z-scale, with warm colors indicating positive values and cold colors indicating negative values.

#### Relative abundance of gut microbiota

Sixty-four bacterial taxa (N = 64) were retained after applying a cutoff retaining only taxa present in more than 70% of the samples, as described in the Methods. Most of the bacteria were identified at the genus level, a few only at higher taxonomic level, while others remained as Unclassified Genus (UCG). See **Fig. 2** for bacteria included, excluded, and their abundance.

**Fig. 2:**
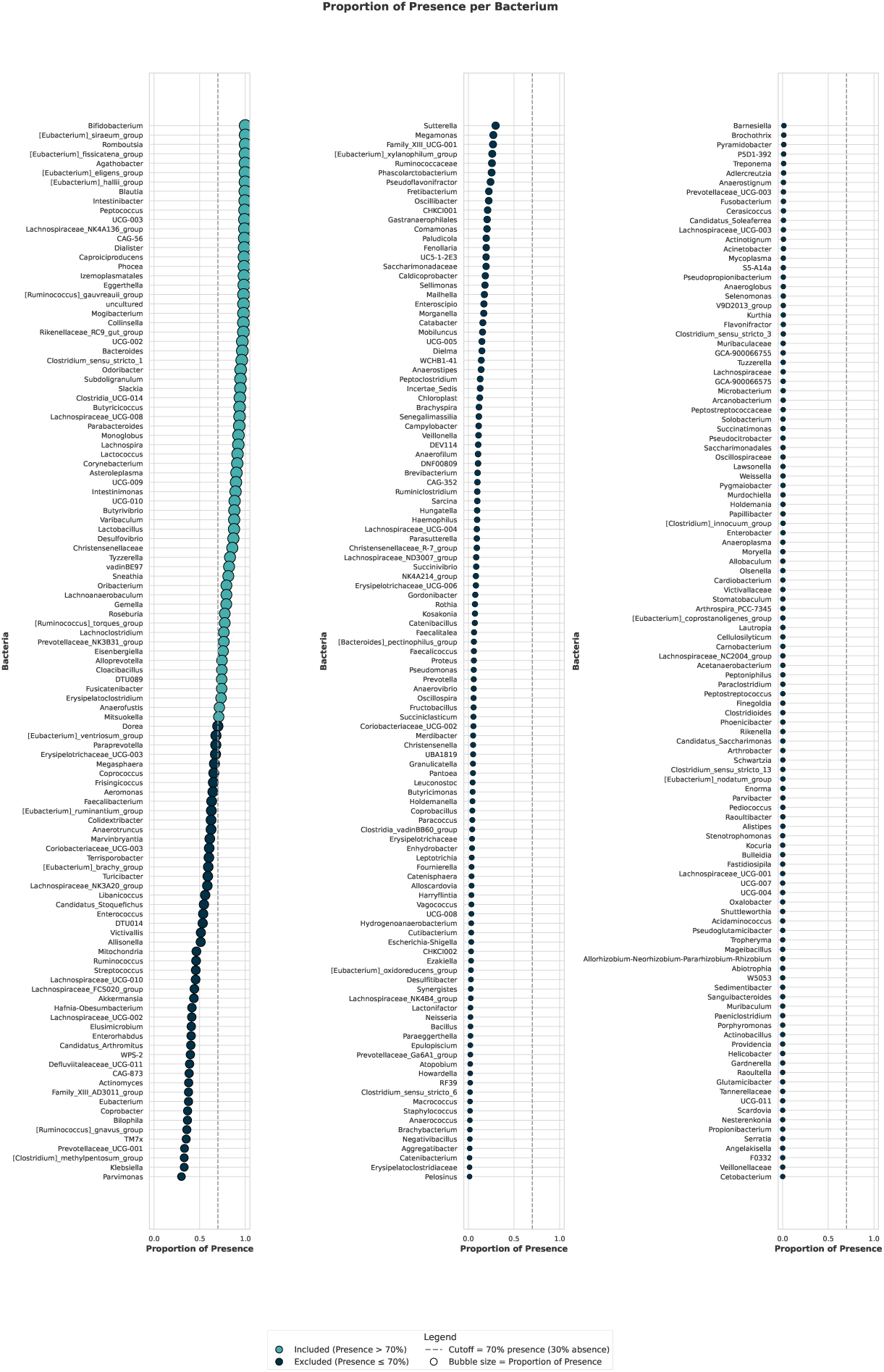
The plot shows the relative abundance of gut microbiota. The x-axis (0–1) indicates the proportion of samples in which each bacteria was detected. A prevalence threshold of *≥* 70% (i.e., *≤* 30% zero values) was applied; retained bacteria are highlighted in light blue. The plot is divided into three panels for readability due to the large number of bacteria identified.

### 2.3 Non-Imaging-Derived Phenotypes (nIDPs)

The non-imaging-derived phenotypes comprised the preregistered stress-, affective, and physiological-related outcome variables intended to supervise the multimodal decomposition. The stress-related measures included physiological markers (cortisol levels and heart rate [HR]), subjective stress ratings, and carotid dilation assessed in response to a classical Socially Evaluated Cold-Pressure Task, SECPT [29]. Prior to the multimodal decomposition, all preregistered nIDPs were screened. For the SECPT-related nIDPs, this screening specifically involved evaluating their validity as indicators of stress induction. As described in the SECPT manipulation-check section and reported in Appendix A, there was a lack of group-level stress response to the experimental manipulation. Consequently, and in deviation from the preregistered analysis plan (see: deviation from preregistration in the Methods section), cortisol, HR, subjective stress, and carotid dilation were excluded from the multimodal decomposition. The remaining twelve nIDPs (N = 12) were used to supervise the multimodal decomposition (see **Fig. 3**). Inspection of their distributions indicated approximate normality for all retained variables except for the CTQ. Although SuperBigFLICA should tolerate some deviations from normality, a log-transformation was applied to the CTQ to improve its distributional properties prior to analysis.

**Fig. 3:**
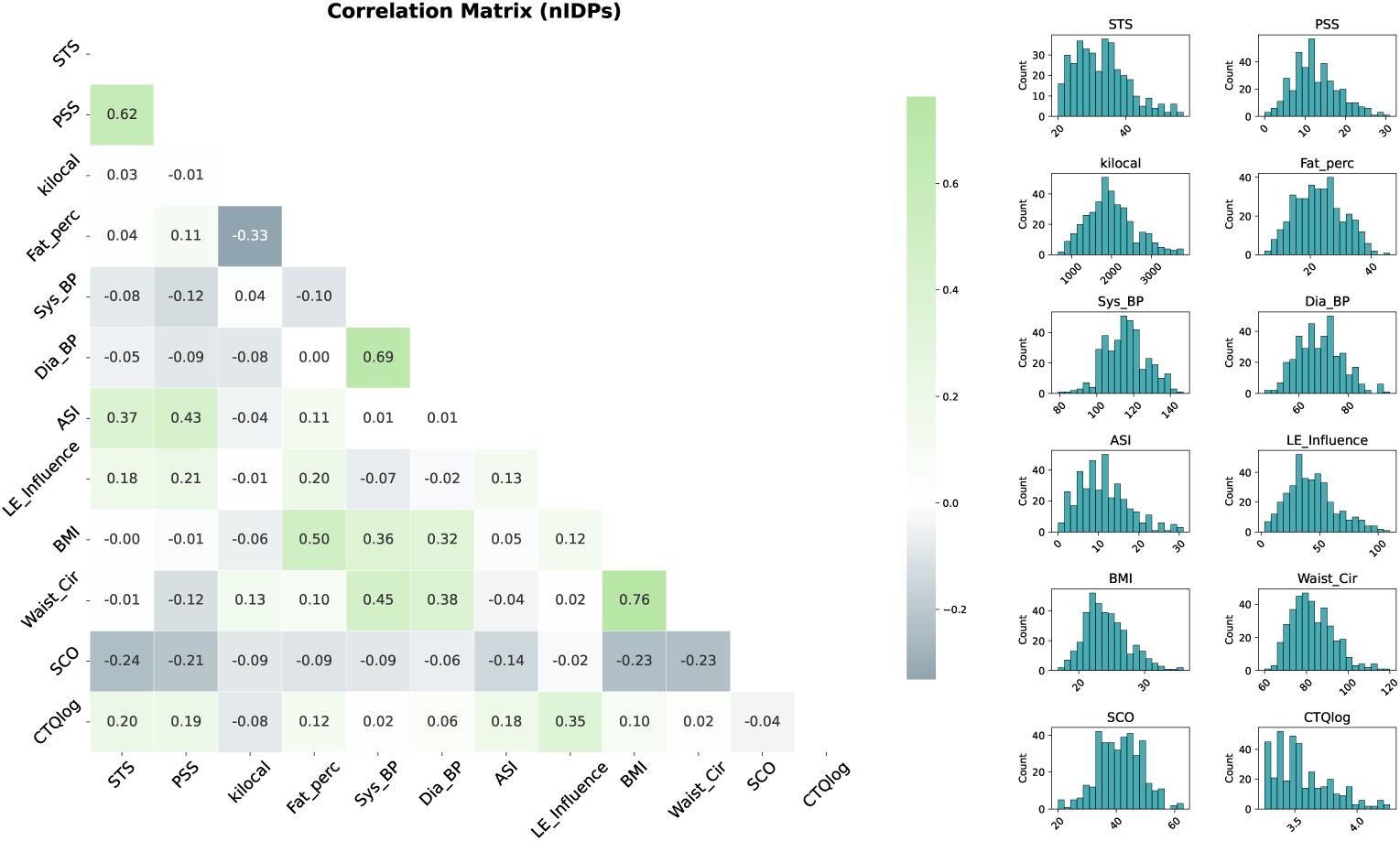
**Features of the selected nIDPs**, including their correlations (left panel) and distributions (right panel). The left panel shows a correlation matrix across all nIDPs: colors toward green indicate positive correlations, colors toward white indicate correlations near zero, and colors toward blue indicate negative correlations. The right panel shows histograms of each nIDP, with counts on the y-axis, illustrating the distribution of values for each variable.

### 2.4 Model selection following hyperparameter tuning

To identify the most robust multimodal decomposition, we first conducted an extensive hyperparameter tuning phase (see Methods). This exploration varied the number of latent components, learning rates, reconstruction weights for IDPs and nIDPs, and sparsity penalties, resulting in a total of 4270 models covering a wide range of plausible parameter combinations. From this large set, models were subsequently filtered based on pre-defined quality criteria (see Methods). Several models survived these cut-offs, providing the basis for further analyses and evaluation of model stability (see **Fig. 4**).

**Fig. 4:**
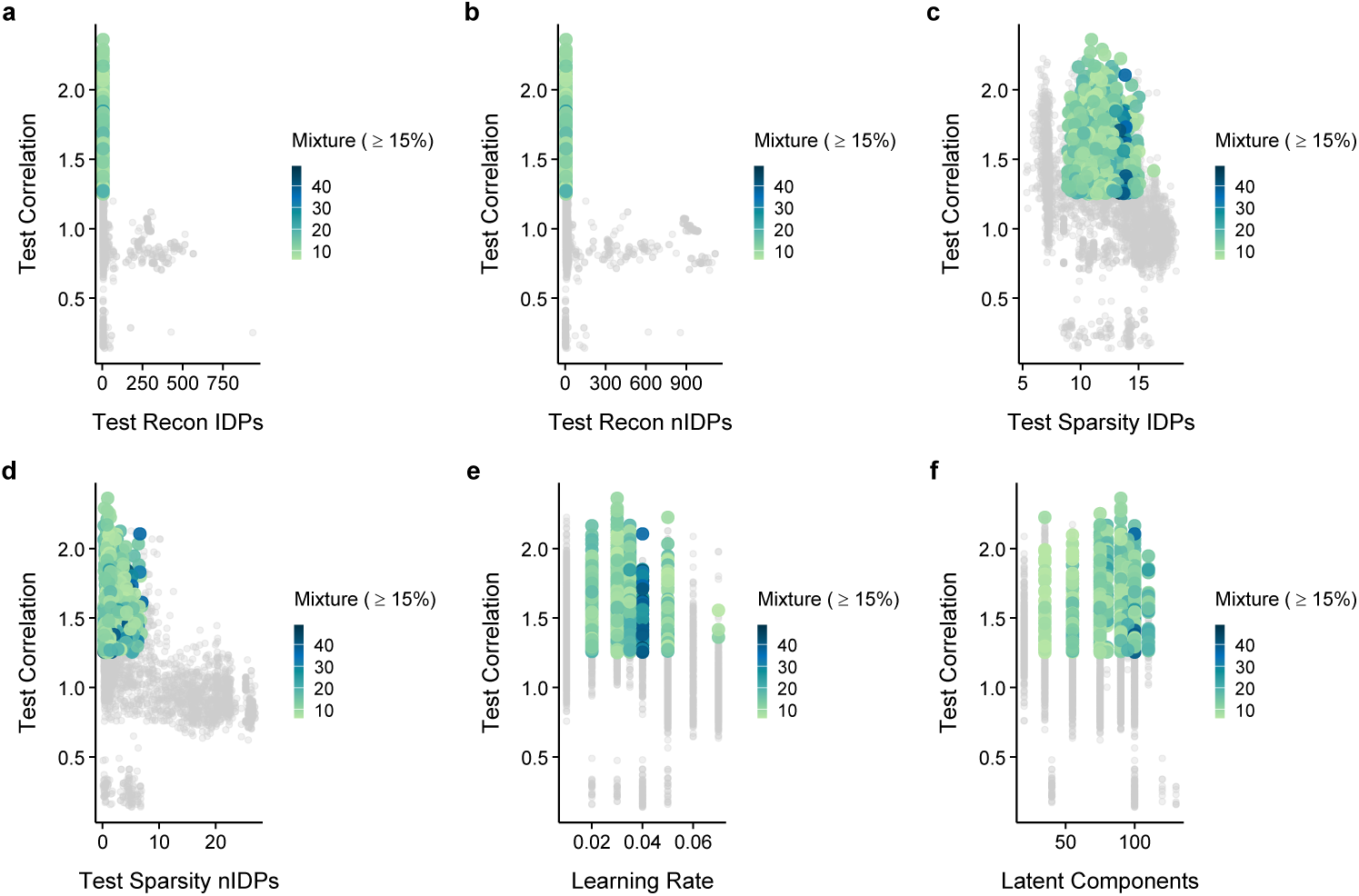
Models before and after filtering. The figure shows the selected models defined as those with a least 15% of mixture between brain and gut-related information. These models are plotted relative to the total correlation of the nIDPs obtained from the test dataset used for validation and: the error of the reconstruction of the IDPs (first lambda), panel **a)**; error of the reconstruction of the nIDPs (third lambda), panel **b)**; the error of the sparsity of the IDPs (second lambda), panel **c)**; the error of the sparsity of the nIDPs (fourth lambda), panel **d)**; the learning rate, panel **e)**; and the number of latent components used, panel **f)**. The models indicated in color gray color were not selected because no brain activations and/or a poor number of components with mixture were found. Selected models are shown in green to blue, with lighter green indicating models with few mixture components and darker blue indicating models with up to 50 mixture components.

The filtered models were then clustered based on all the standardized parameters used in (and errors produced by) the SuperBigFLICA algorithm from the test dataset. From this clustering process, five clusters of models were obtained (see **Fig. 5**). Within each cluster, the centroid model was selected in a subsequent, independent General Linear Model (GLM) analysis.

**Fig. 5:**
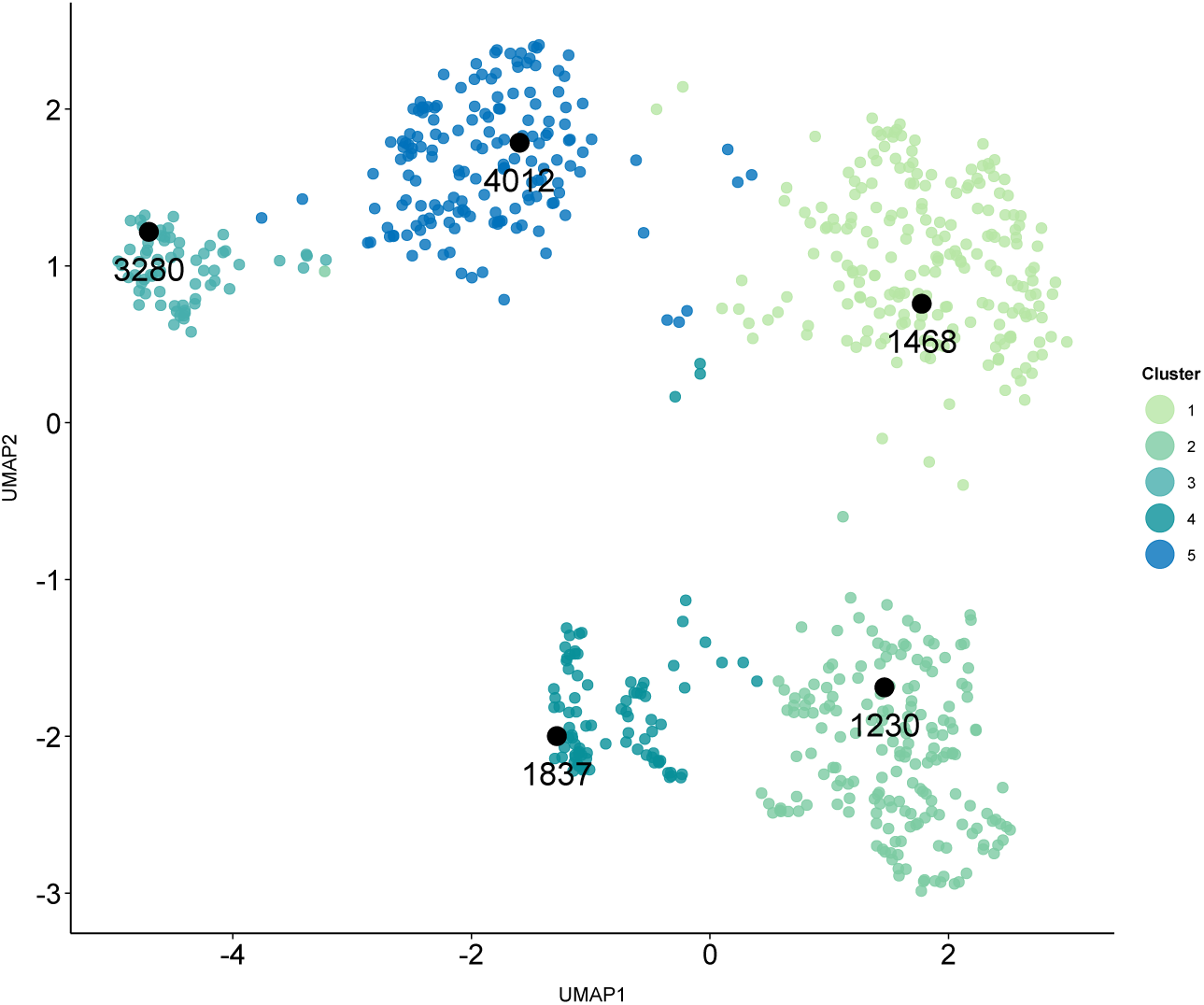
Display of the five clusters (and the centroids) obtained after filtering procedures. Model 4012 included 12 components with mixture, while models 1468, 1230, 1837 included 13 components with mixture each. Model 3280 included 28 components with gut-brain mixture (Comp1, Comp15, Comp23, Comp25, Comp26, Comp27, Comp28, Comp30, Comp35, Comp40, Comp47, Comp49, Comp52, Comp59, Comp60, Comp64, Comp65, Comp67, Comp75, Comp77, Comp80, Comp83, Comp90, Comp93, Comp94, Comp96, Comp97, Comp99).

### 2.5 Prediction of the behavioural data

General Linear Models were used to explore if and which of the selected models significantly predicted behavioural outcomes from the food auction task. It is important to highlight that these data were not included in the model discovery and selection phase. Among the 5 representative models, model number 3280 was the one with the highest number of mixture components (N = 28 of 100). Furthermore, one of these mixtures, component 93, showed the clearest associations with some nIDPs and was predictive of behaviour. See **Fig. 6** for the model summary. Specifically, component 93 was characterized by a mixture of LN (17.3%), DMN (11.1%) and gut microbiome (71.3%) with several bacteria above the threshold ((*z >* 2.3) or (*z <* 2.3)) including UCG-010, *z* = 2.90 and DTU089, *z* = 3.12; this component loaded high especially in BMI (1.81), ASI (0.36), and life event influence (0.73), and was predictive of both valuation (*β* = 0.17, SE = 0.08, *t* = 2.23, *p*-value = 0.026) and choice (*β* = 0.16, SE = 0.06, *t* = 2.58, *p*-value = 0.010) of unhealthy (vs. healthy) food items. See **Fig. 7** for specifics about component 93 including clusters equal to-or larger than 20 voxels in Table B1 and Table B2 of the supplementary information in Appendix B.

**Fig. 6:**
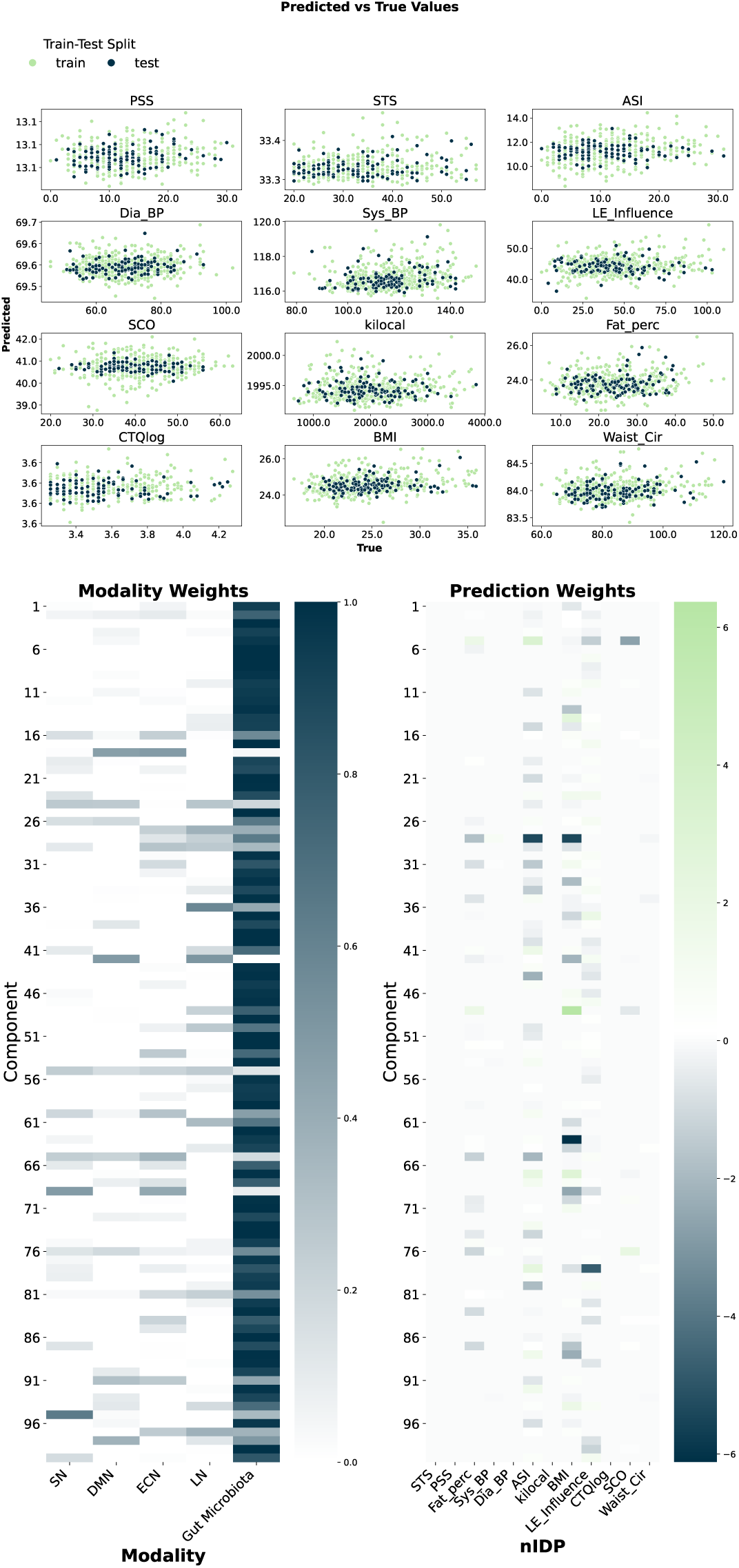
Summary of the predictive model (3280). The top panel visualizes the model fit (predicted vs. true values) for each nIDP from the test (blue) vs the training (green) dataset (total of 12 models fits).^1^T^0^he panel on the left-bottom displays the heat maps of the modality weights for each component of each modality input; while the panel on the right-bottom displays the prediction weights for each component of each nIDP.

**Fig. 7:**
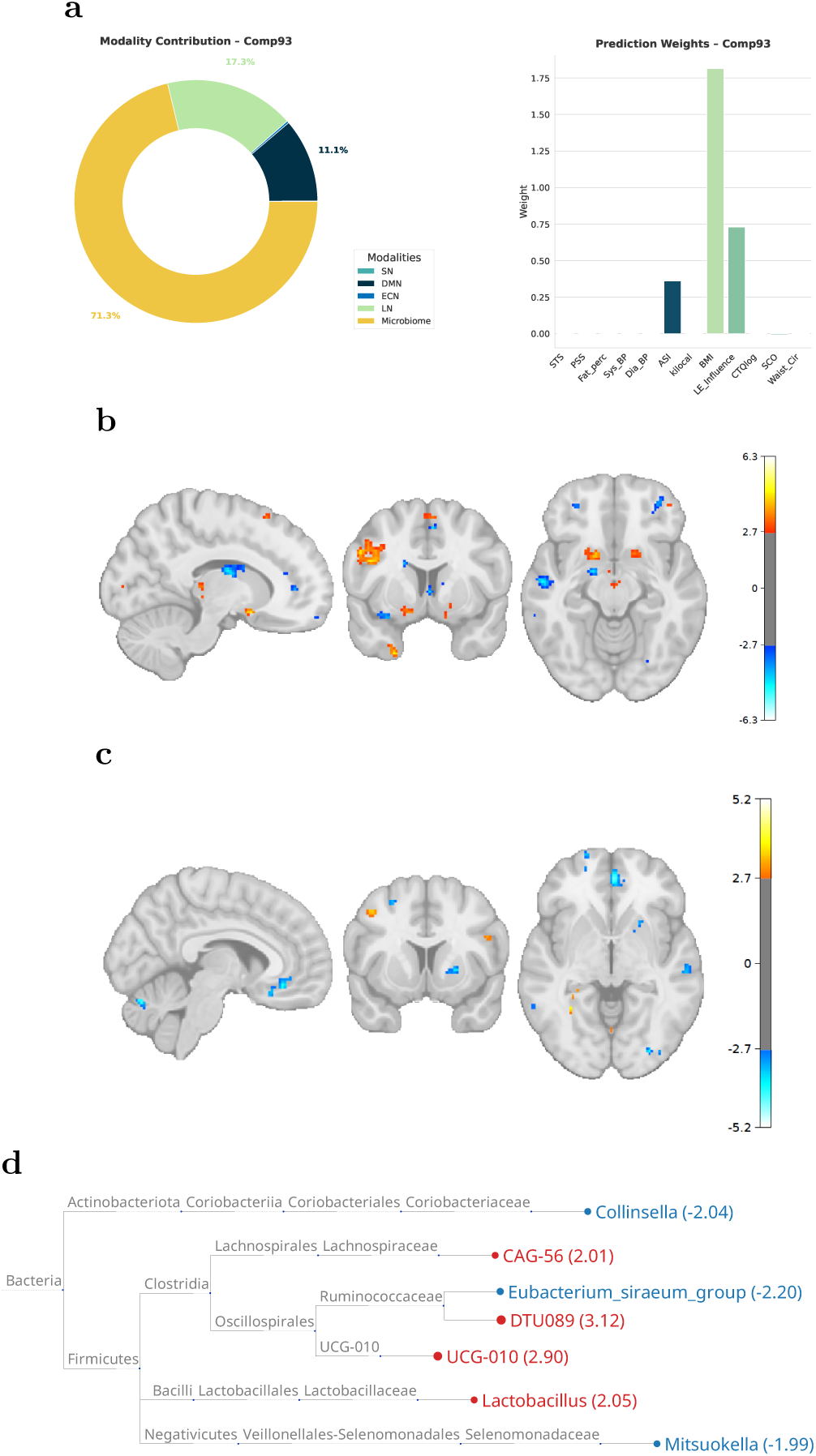
Specific features of component 93 of model 3280. **a)** Radar plot of the percentage modality contribution to Component 93 (LN 17.3%, DMN 11.1%, gut microbiota 71.1%), and bar plot of the component’s standardized associations with the nIDPs (strongest loadings: BMI = 1.81, life-event influence = 0.73, ASI = 0.36). **b)** Significant activation map for the LN, with a zoom into the NAc (*x* = *−*12, *y* = 20, *z* = *−*14) and dorsal striatum. **c)** Significant activation map for the DMN. Maps in b) and c) are thresholded at *|z|* = 3 with a minimum cluster size of 20 voxels. **d)** Phylogenetic tree of taxa above (*|z| >* 2.3, red) and below the threshold; UCG-010 (*z* = 2.90) and DTU089 (*z* = 3.11) exceeded the predefined threshold used in [19].

**Fig. 8:**
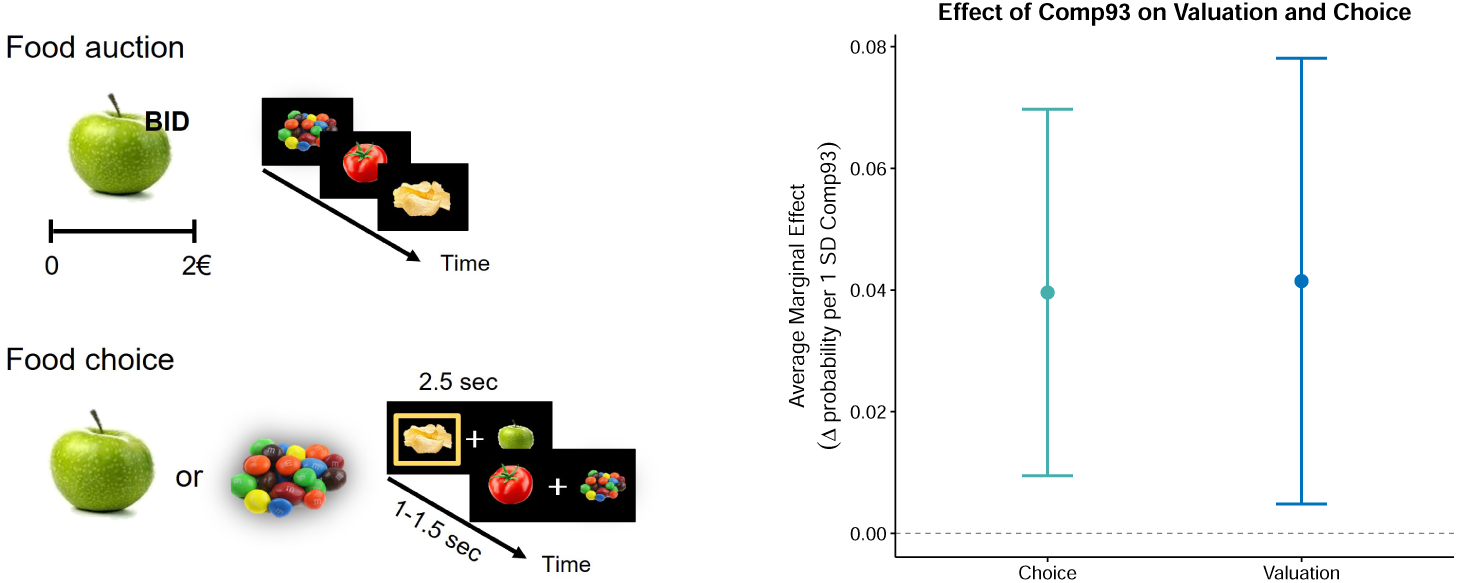
The food auction and choice task is displayed on the left. During this task, participants first placed monetary bids for individual food items (food auction) to reveal subjective value. These bids were then used to construct pairs of healthy and unhealthy foods with approximately equal subjective value. Participants subsequently chose between these pairs. The plot on the right shows the average marginal effect (AME) of a 1-SD increase in the Component 93 loading on the probability of valuing and choosing unhealthy over healthy food items, estimated from the GLM adjusted for all 100 latent components; error bars denote 95% confidence intervals. A 1-SD increase was associated with an approximately 4% higher probability of selecting unhealthy items (choice: *β* = 0.16, SE = 0.06, *t* = 2.58, *p* = 0.010; valuation: *β* = 0.17, SE = 0.08, *t* = 2.23, *p* = 0.026). Estimates were derived from held-out food auction task data not used in model training.

Descriptively, follow-up analysis of the average marginal effect (AME) revealed that after adjusting for all 100 components, a 1-standard-deviation increase in this component was associated with an approximately 4% higher probability of selecting unhealthy food items, see section 2.5. The association of component 93 with behaviour remained significant after adjusting for BMI (all p-values below 0.05), indicating that its association with unhealthy food valuation and choice reflects eating behaviour-relevant variance in the gut–brain configuration rather than a simple reflection of body mass index. Restricting the analysis to each participant’s top-3 most-valued items per food category yielded consistent results with a small increase in effect size (Valuation: AME = increased from 0.041 to 0.042; Choice: AME = increased from 0.039 to 0.044). Furthermore, we evaluated whether the association of this component was influenced by gender by including gender as a covariate in the GLM model. Gender did not alter the association of component 93 for either valuation (*p*-value = 0.026) or choice (*p*-value = 0.011) of unhealthy versus healthy foods. We also tested whether this component, adjusted for all other components, was associated with sugar intake as measured by the FFQ. This analysis was not significant (*p*-value = 0.41).

For transparency and to increase understanding of this model, we also explored a component that was associated with behaviour but did not show a substantial mixture (Component 70). This component was negatively predictive of valuation (*β* = - 0.12, SE = 0.05, *t* =-2.12, *p*-value = 0.035) and choice (*β* =-0.10, SE = 0.04, *t* =-2.12, *p*-value = 0.034) of unhealthy food and included only the gut microbiota modality in combination with elevated BMI. Component 70 contained taxa with substantially higher absolute z-values than Component 93. First five positive taxa: Roseburia (*z* = 8.26), Lactobacillus (*z* = 5.58), Lachnospiraceae UCG-008 (*z* = 5.39), Collinsella (*z* = 5.26), Cloacibacillus (*z* = 4.84). First five negative taxa: [Eubacterium] eligens group (*z* = *−*5.13), [Eubacterium] hallii group (*z* = *−*5.48), Slackia (*z* = *−*5.75), Lachnospiraceae NK4A136 group (*z* = *−*6.50), [Eubacterium] siraeum group (*z* = *−*6.80). See the full list of taxa in Appendix C. However, analysis on the average marginal effect revealed that after adjusting for all 100 components, an increase in standard deviation of 1 in this component was associated with an almost 0 probability of selecting unhealthy food items. Moreover, when BMI was included as a covariate in the GLM, this component was no longer significant, suggesting that its association with unhealthy-food choice was largely accounted for by BMI rather than by the component itself. This was not the case for Component 93, whose association remained significant after BMI adjustment.

### 2.6 Follow-up checks of model robustness

As a final step, we conducted several follow-up checks. First, to ensure that the train and test data sets used were statistically comparable in their nIDPs and confounders, we checked for potential differences in their distributions (train-test distribution check). The results of the *t*-tests did not reveal significant differences (all *p*-values above 0.05), see Appendix D.

Second, we evaluated the stability of the supervised FLICA model with respect to random initialization. Although the most frequent correlation values point to a mild correlation (range between 0.1 and 0.3 across the nIDPs), these values are in line with those reported within the original paper in which SuperBigFLICA was developed [18].

Third, we evaluated the stability of the supervised FLICA model with respect to random train–test splits. Across the hundred models trained, we observed a frequent mild correlation between predicted and true values of the ASI, BMI, and life events influence, see **Fig. 9b**. This highlights the sensitivity of model performance to data partitioning.

**Fig. 9:**
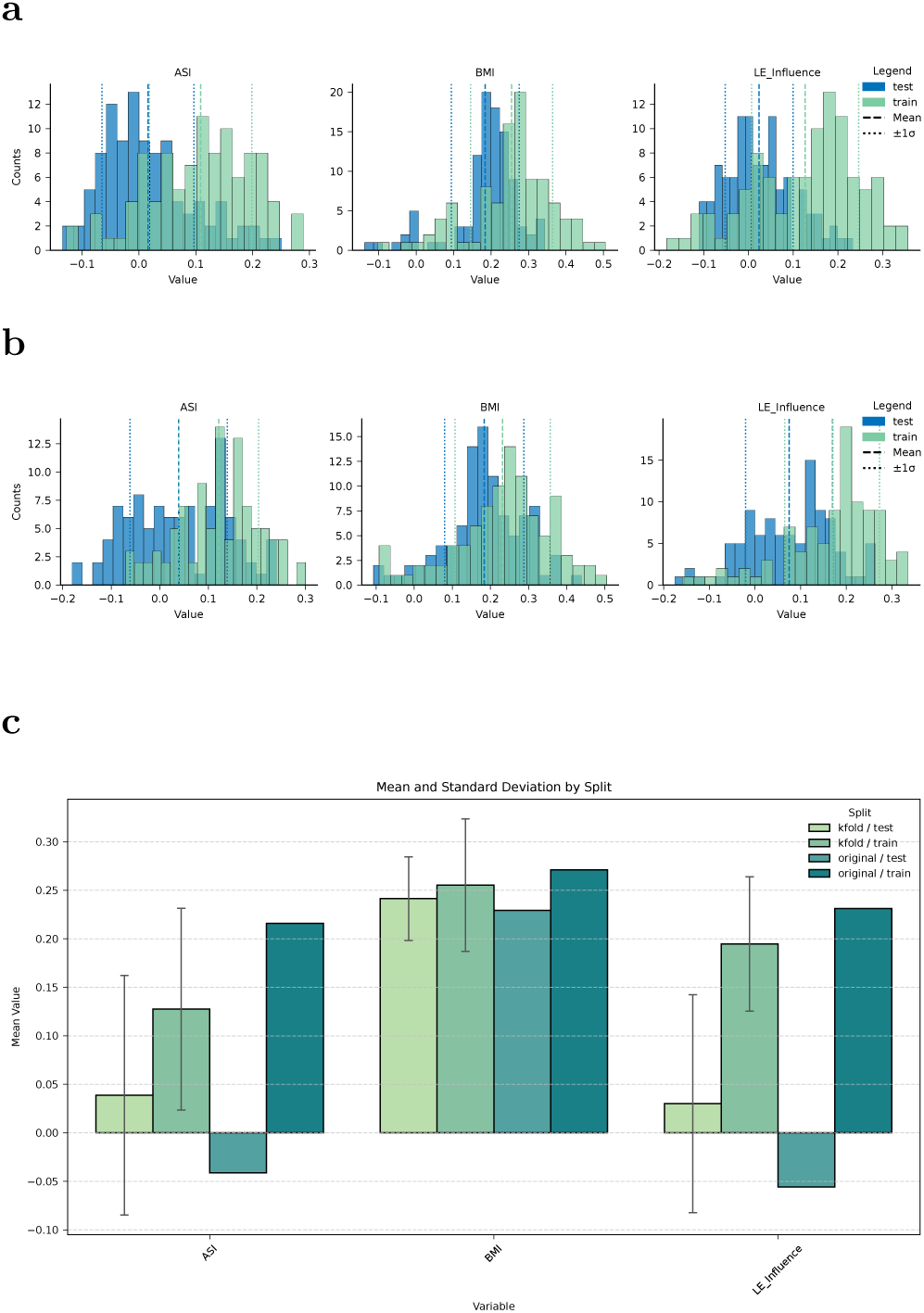
Robustness of the supervised FLICA model across three stability checks. Distributions of Pearson correlations between predicted and observed nIDP values (test set, blue; training set, green) for the three most strongly loading nIDPs (ASI, BMI, life-event influence). (**a**) Counts of correlations across 100 random initializations. (**b**) Counts of correlations across 100 random train–test splits. (**c**) Correlations from 5-fold cross-validation (fold-averaged, shown against the original fold) for ASI, BMI, and life-event influence. Across checks, test-set correlations were predominantly in the mild range (median *r* between 0.1 and 0.3), consistent with values reported for SuperBigFLICA in [18]. Model 3280 parameters were held constant (100 components, learning rate = 0.04, 300 iterations; *λ* = 0.75, 0.2, 0.6, 0.2); *n* = 592.

Fourth, a 5-fold cross-validation was performed to further assess the stability and generalizability of the model across different data partitions. The resulting correlations were averaged across folds and plotted alongside those obtained from the original fold for ASI, BMI, and life events influence, see **Fig. 9c**. Among the observed correlations, the one that involved BMI remained consistent with the patterns reported in the original work that described the development of the algorithm. The correlation of ASI and life events remained less consistent.

Finally, we checked whether component 93 reappeared in similar models included in cluster 3. Within Cluster 3, component 93 repeatedly reappeared in many models, see table 1, indicating a reproducible signal. Most models showed similar patterns of connectivity and gut microbiota associations.

**Table 1:** Models within cluster 3 in which component 93 was present and significantly associated with behaviour. Of note, all the nIDPs refer to a positive increase.

| Model ID | Modality | Estimate | features (nIDPs) |
| --- | --- | --- | --- |
| 3344 | DMN (7.8%), LN (12.8%), Gut (71.3%) | −0.14 | ASI, BMI, LE_influence |
| 3304 | DMN (32.7%), LN (11.7%), Gut (55.6%) | 0.16 | ASI, BMI, LE_influence |
| 3297 | DMN (27.8%), LN (19%), Gut (53.1%) | 0.13 | ASI, BMI, LE_influence |
| 3272 | DMN (36.7%), LN (29.2%), Gut (33.3%) | 0.13 | ASI, BMI, LE_influence |
| 3285 | DMN (27.7%), LN (22.7%), Gut (49%) | 0.14 | ASI, BMI, LE_influence |
| 3232 | DMN (39.6%), LN (20.4%), Gut (39.5%) | 0.15 | ASI, BMI, LE_influence |
| 3298 | DMN (26.5%), LN (19%), Gut (54.4%) | 0.14 | Predominantly BMI |
| 3281 | DMN (13.1%), LN (17.8%), Gut (68.7%) | 0.16 | BMI |
| 3286 | DMN (29.6%), LN (23.2%), Gut (46.6%) | 0.13 | BMI |
| 3275 | DMN (19.6%), LN (9.5%), ECN (23.6%), Gut (47%) | 0.13 | BMI, LE_influence, SCO |
| 3263 | DMN (5.2%), LN (8.8%), ECN (28.9%), Gut (57.7%) | 0.12 | BMI, LE_influence, SCO |
| 3303 | DMN (34.2%), LN (14.5%), Gut (51.2%) | 0.18 | ASI, BMI, LE_influence, SCO |
| 3292 | DMN (32.3%), LN (7.4%), Gut (60.1%) | 0.12 | ASI, BMI, LE_influence, SCO |

## 3 Discussion

This preregistered study had a twofold aim: to provide a proof-of-concept for our integrative approach, and to test whether elevated valuation and choice of unhealthy food can be predicted by latent, multivariate associations between reward-related brain networks and gut microbiota composition, guided by stress-and affect-related individual differences. Applying supervised linked independent component analysis (SuperBigFLICA) in a large, non-clinical cohort (N = 592), we established proof-of-concept by recovering a candidate phenotype consistent with our hypothesis: a latent component jointly characterizing a specific gut microbial profile, resting-state connectivity in limbic and default-mode regions, and elevated anxiety sensitivity, life-event impact, and body mass index. This component predicted elevated valuation and choice of unhealthy food in the held-out food auction task, albeit with a small effect size.

Among the five representative models retained after clustering, model 3280 was distinguished by its capacity to capture a large number of components with a meaningful gut–brain mixture, that is, components reflecting coordinated variation across both brain connectivity and gut microbiota, rather than either modality in isolation. Within this model, component 93 stood out as the only component combining this gut–brain mixture with clear associations with some of the psychological and physiological nIDPs. It was characterized by 71% gut microbiota contribution, 17% Limbic Network, and 11% Default Mode Network. It loaded most strongly on BMI, followed by the impact of life events and anxiety sensitivity, a constellation that characterizes individuals who are not clinically ill but who might carry a heavier affective burden in daily life. This same component was associated with both food valuation (how much participants were willing to pay for unhealthy items) and food choice (whether they selected unhealthy over healthy items when value was held constant), two measures that were strongly correlated in this sample (Spearman r = 0.83). The results from this exploratory study suggest that gut-brain linkage in this latent component may reflect a general disposition toward unhealthy food and should be further examined in confirmatory studies.

Within the Limbic Network, component 93 was characterized by increased connectivity involving the NAc, together with relatively reduced engagement of the caudate, amygdala, and parahippocampal gyrus connectivity. The NAc is a central node of the mesolimbic dopamine system and has been extensively implicated in the assignment of incentive salience to rewarding stimuli, including food [30]. In the context of food intake specifically, NAc activity tracks the motivational value of palatable foods and is sensitive to dopaminergic modulation [31, 32]. Alterations in NAc have been linked to emotional eating behaviours and to mediate relationships between these behaviours and adiposity [33]. Additionally, in humans under stress, emotional eaters show disrupted NAc responses during food reward anticipation, with altered NAc activation specifically associated with stress-induced changes in eating [34]. Furthermore, reduced caudate engagement might reflect a shift away from goal-directed action-outcome processing toward habitual stimulus-response control [35]. In the food domain, this could translate into a stronger tendency to respond to the hedonic (unhealthy) qualities of food rather than to longer-term health consequences. Additionally, decreased amygdala and parahippocampal coupling has been associated with disrupted encoding of affective and contextual memory traces that link food cues to their anticipated consequences, a process that may be particularly impaired in the context of elevated anxiety sensitivity and life-event burden characterizing this component. This is consistent with broader evidence that anxiety and stress alter the affective weighting of food-related stimuli, biasing responses toward immediate palatability [5].

Within the DMN, reduced connectivity of paracingulate and prefrontal nodes is known to exist in individuals with elevated binge-eating tendencies [36, 37], and has been interpreted as reflecting less efficient intrinsic organization of circuits involved in self-regulatory processing. The DMN, and especially the medial prefrontal cortex, plays a key role in internally directed thought, including the anticipation of future outcomes and self-referential decision-making [38]. In the context of dietary choices, prefrontal engagement has been associated with successful dietary self-regulation through modulation of ventromedial Prefrontal Cortex (vmPFC) value signals [39]. Accordingly, the connectivity pattern observed in the DMN of component 93 may reflect a weaker baseline organization of the circuits that support the integration of health goals into food valuation, a disposition that may predispose individuals toward hedonic choices particularly when cognitive resources are taxed by affect or life stress. Moreover, two further features of Component 93 deserve a brief mention: the putamen and subgenual Anterior Cingulate Cortex showed opposite polarities across the LN and DMN; and the insula showed a similar dissociation, with decreased left posterior-inferior insula coupling within the LN and increased right mid-insula coupling within the DMN. These regional patterns reflect relative associations of the regions to the two networks, which while opposed do not necessarily imply uniform regional effects. A more specific interpretation, particularly for the insula subregions, which differ in their roles in interoceptive processing [40], would require task-based data. In sum, the resting-state connectivity pattern of component 93 resembles what has been observed in individuals with elevated BMI and heightened emotional reactivity to food [36, 37, 41]: a phenotype marked by enhanced intrinsic engagement of incentive-salience circuits and reduced caudate-PFC involvement.

Component 93 also involved a specific gut microbiota configuration, characterized by a high relative abundance of UCG-010 and DTU089, two poorly characterized, uncultured genera within the class *Clostridia*. DTU089 has been described almost exclusively in relation to host metabolic and skeletal phenotypes: it is more abundant in individuals with lower dietary protein intake [42] and has been negatively associated with bone density [43]. To our knowledge, no study has linked it to brain function, eating behaviour, or affect, and its role in the present component therefore remains to be determined. One tentative interpretive avenue is metabolic: several fibre-fermenting *Clostridia* contribute to microbial tryptophan metabolism via the indole pathway, which yields derivatives such as indole-3-acetic acid (IAA) [44]. Direct evidence that DTU089 produces IAA is lacking, and the capacity to generate indole derivatives such as IAA is restricted to particular bacterial species rather than being a general feature of the gut microbiota [45]; this pathway is nonetheless of interest here because of a partial convergence with Osadchiy and colleagues [46]. In sixty-three healthy adults, they related faecal concentrations of indole, IAA, and skatole to resting-state functional and structural connectivity of the amygdala, NAc, and anterior insula, and to BMI, food-addiction scores, and anxiety. These phenotypic and neural features partially overlap with the nIDPs structuring our decomposition (BMI and anxiety sensitivity) and with the regions defining the LN–DMN pattern of Component 93, most notably the NAc. The co-occurrence of elevated DTU089 and altered NAc connectivity within a single latent component is thus consistent with (though far from establishing) a tryptophan–indole-mediated gut-to-brain route relevant to unhealthy food intake.

Less can be said about UCG-010. Functional characterization of this genus in humans is very limited, and the available evidence associates *Oscillospiraceae*, including UCG-type genera, with leaner phenotypes: reduced diversity within this family is broadly linked to higher BMI [47]. In the present component, however, UCG-010 loaded positively alongside elevated BMI, a direction that runs counter to this general trend. Given how little is established about this uncultured taxon, we do not over-interpret its contribution, which may partly reflect the multivariate, network-level structure of the decomposition rather than a direct association with adiposity; its role in this phenotype remains to be clarified.

In sum, while direct mechanistic conclusions cannot be drawn in the absence of metabolomic data, the convergence with Osadchiy et al. supports the biological plausibility of an altered balance of microbial tryptophan-metabolite production modulating NAc connectivity, a hypothesis best tested through targeted metabolomics and mechanistic investigation.

It should be noted that, while the absolute effect size of component 93 was modest (a 4% higher probability of selecting unhealthy food items per 1 SD increase in component 93 score), this estimate represents the marginal contribution of component 93 over and above 99 other latent components, and is therefore conservative. The magnitude of this effect is difficult to relate to comparable studies in this domain, since to our knowledge no previous study in food decision-making has applied a multi-component ICA decomposition jointly across brain and microbiome data in healthy humans. Osadchiy et al. [46], whose findings most directly parallel the present result at the metabolite level, reported bivariate Spearman correlations in the range of *r ≈* 0.31–0.60 between fecal indole metabolites and resting-state functional connectivity of the NAc and amygdala, and between those connectivity measures and food addiction scores (YFAS), in a sample of 63 healthy adults. Those associations were unadjusted for the broader variance structure of the brain or microbiome data, and the YFAS is a questionnaire-based measure of food addiction tendency rather than a direct behavioural index of choice. By contrast, the effect reported here derives from a marginal estimate adjusted for the full 100-component model and is obtained from a behavioural task rather than self-report. In this context, a marginal effect of approximately 4% per 1 SD is arguably relevant at the (healthy) population level.

In contrast to Component 93, where the co-occurrence of microbial, neural, and behavioural features pointed toward a coordinated gut–brain phenotype linked to incentive-driven food valuation and choice, the configuration of Component 70 is more parsimoniously interpreted as a BMI-coupled microbiome signature potentially shaped by dietary patterns rather than a reward-driven pathway. The taxa loading on this component are functionally heterogeneous but converge on a fermentation-related ecological niche, encompassing fibre degradation, short-chain fatty acid (SCFA) production, and cross-feeding interactions [48–50]. A proportional weakening of this integrated network would be expected to subtly reduce SCFAs and other gut-derived signalling molecules [44, 50, 51]. The behavioural pattern associated with Component 70 (an inverse association with unhealthy food choices, in the absence of a neural signature, and tightly coupled to elevated BMI) differs qualitatively from the reward-driven profile of Component 93. One speculative possibility is that Component 70 reflects a restriction-related rather than a value-driven route to food choice in the food auction task: higher-BMI participants may be more likely to engage in active dietary restraint, which could in principle shape microbiota composition through reduced intake of fermentable substrates while also influencing food-choice behaviour. If this were the case, Components 93 and 70 might capture two distinct routes that have long been hypothesized to coexist but are rarely dissociated in the same model: a value-based, reward-driven route in which incentive salience and weakened self-regulatory organization jointly bias choices toward palatable items, and a restriction-related, BMI-coupled route in which their microbiota correlates accompany dietary patterns. Although this interpretation is plausible, we emphasize that it is speculative. Disentangling these possibilities will require longitudinal designs combined with explicit measures of dietary restraint, eating-related cognitions, and current dieting behaviour, alongside microbiota profiling, peripheral metabolite assessment, and task-based neural measures of food valuation.

In sum, the results of this study support the view that gut–brain interactions relevant to food valuation and choice are inherently multivariate and are unlikely to be fully resolved through unidimensional analyses of either modality. Importantly, this sample of participants was non-clinical, with no diagnosed eating disorders, suggesting that this phenotype may represent a subclinical predisposition detectable through multivariate analysis.

Several limitations should be considered when interpreting the findings of this exploratory study. First, two features of the sample constrain the strength of the conclusions. Although large (N = 592) relative to many multimodal neuroimaging studies, it remains modest compared to the dataset used to develop the SuperBigFLICA algorithm, which likely contributes to the sensitivity of model performance to data splits and initialization and highlights the data-hungry nature of high-dimensional supervised fusion methods. Beyond size, the cohort was not designed with eating behaviour or overeating in mind; as a general-population sample, it captures only limited variation in the phenotypes most central to our hypothesis. Replication is therefore needed not only in larger cohorts, but specifically in datasets characterised for eating-related traits, in which the hypothesised gut–brain contribution to unhealthy-food valuation can be tested more directly.

Second, all neural measures were derived from resting-state functional connectivity. While this approach is well-suited for identifying stable network configurations [52], it does not permit direct inference about task-evoked reward processing or causal mechanisms. This might also explain the relatively small effect size in predicting behaviour, as we did not measure fMRI connectivity during the behavioural task. Future work combining gut-related information with resting-state and task-based paradigms will be essential to clarify how these intrinsic network biases translate into food-related decision processes.

Third, the cross-sectional nature of this study precludes conclusions about directionality or causality. It remains unclear whether the identified microbial and neural features contribute to elevated unhealthy food valuation/choice, emerge as a consequence of dietary patterns, or reflect reciprocal interactions over time.

Fourth, the absence of a reliable acute stress response represents an important limitation, particularly given the use of a healthy, non-clinical population. In the present samples, the inclusion of valid acute stress-reactivity markers as nIDPs would have been highly informative, potentially enabling a more precise decomposition of latent brain–gut components associated with stress-induced changes in food valuation. Finally, behavioural relevance was established for unhealthy-food valuation and choice specifically; whether this phenotype generalizes to other behavioural domains was not tested here. Future longitudinal and interventional studies will therefore be needed to test its predictive validity across behaviours, contexts, and populations.

In conclusion, this study makes two complementary contributions. First, it demonstrates the feasibility of supervised gut–brain integration for recovering behaviourally relevant structure: an appropriately trained multivariate model surfaced a candidate phenotype linked to food-choice behaviour, which, to our knowledge, has not previously been shown in this domain. Second, it characterizes a phenotype in which individuals with higher anxiety sensitivity, greater life-event impact, and higher BMI appear more likely to express a configuration marked by enhanced intrinsic reward valuation, diminished regulatory integration, and a distinct microbiota milieu. Even in the absence of acute stress reactivity, this latent configuration may predispose toward hedonic eating when palatable food cues are encountered.

## 4 Methods

The dataset was extracted from the HBS cohort. HBS is a large longitudinal study conducted in Nijmegen, The Netherlands, which involved about 1000 healthy adult volunteers. The study includes three longitudinal assessments distributed equally within a one-year time window. Each assessment includes a variety of measures obtained at home and on campus (i.e., pre-visit online questionnaires, burst week - real-world assessments, lab visit on campus, and post-visit online questionnaires). The Institutional Review Board of Radboud University Medical Center approved the HBS on the 23rd of May, 2019 (reference number: 2018–4894) in accordance with the latest revision of the Declaration of Helsinki [53]. For more information, see the original document [20].

### 4.1 Deviation from the preregistered analysis

First, contrary to what was preregistered, we did not include an independent test dataset. Although we refer to one split as the “test” dataset, this dataset functioned as a validation set used for model selection within the SuperBigFLICA framework. This terminology follows conventions within SuperBigFLICA and should not be interpreted as a classical independent test set. Because of the limited sample size, the entire dataset was employed for model training and validation, and no separate test set was reserved.

Second, although the SCO measure was initially planned for inclusion in the exploratory analyses, we incorporated it as a non-imaging-derived phenotype (nIDP). This was because cognitive control constitutes an inverse marker of stress and its inclusion served as an internal validity check via correlation analyses with the other nIDPs (see correlation matrix, Figure 3).

Third, menstrual cycle information was not included in our exploratory analyses because the index we received did not contain any reported subjective symptoms related to menstrual phases, such as emotional dysregulation or emotional reactivity. Finally, as reward valuation processes also involve the activity of the orbital prefrontal cortex, and decision making processes also involve executive control-related brain areas activations, such as the dorsolateral prefrontal cortex, DMN and ECN were included in addition to the two preregistered networks (i.e., SN and LN).

### 4.2 Data extraction

For the purpose of the present exploratory study, we specifically included: *demography* characteristics (such as gender); *biological* data including anatomical brain data, functional magnetic resonance imaging data during a resting state (rsfMRI), gut microbiota composition (e.g. relative abundance of bacteria genera) obtained from stool sample, salivary cortisol levels in response to the SECPT; *physiological* parameters including HR and carotid dilation in response to the SECPT and blood pressures (diastolic and systolic) at rest; *behavioural* data consisting in the performances obtained during the FAT (which includes a valuation and choice part); *subjective* measures such as subjective stress measured by the Perceived Stress Scale, subjective stress levels perceived in response to the SECPT, scores from the State and Trait Anxiety Inventory, Anxiety Sensitivity Index, Childhood Trauma Questionnaire, (threatening) life events, and cognitive control; *body composition* data such as fat percentage, body mass index, and waist circumference; and *diet-related* information extracted from the Food Frequency Questionnaire such as kilocalories consumed.

As the first assessment included the highest rate of participation in the FAT, we requested to include the measures listed above from this assessment. The inclusion and exclusion criteria included in the present study are those already employed at the time of the execution of the HBS (which can be found here [20]).

Statistical outliers were defined using an Interquartile Range (IQR) of 2 within-subject and applied to: each nIDP, subjective loadings from any component tested in the GLMs and to the behavioural measures (i.e., unhealthy vs. healthy food ratios, one for valuation and the other for choice)

### 4.3 Analysis Pipeline

Our analysis followed a multi-stage pipeline designed to extract stable and behaviourally predictive latent components from multimodal data integrating four brain networks and gut microbiota, see **Fig. 10**.

**Fig. 10:**
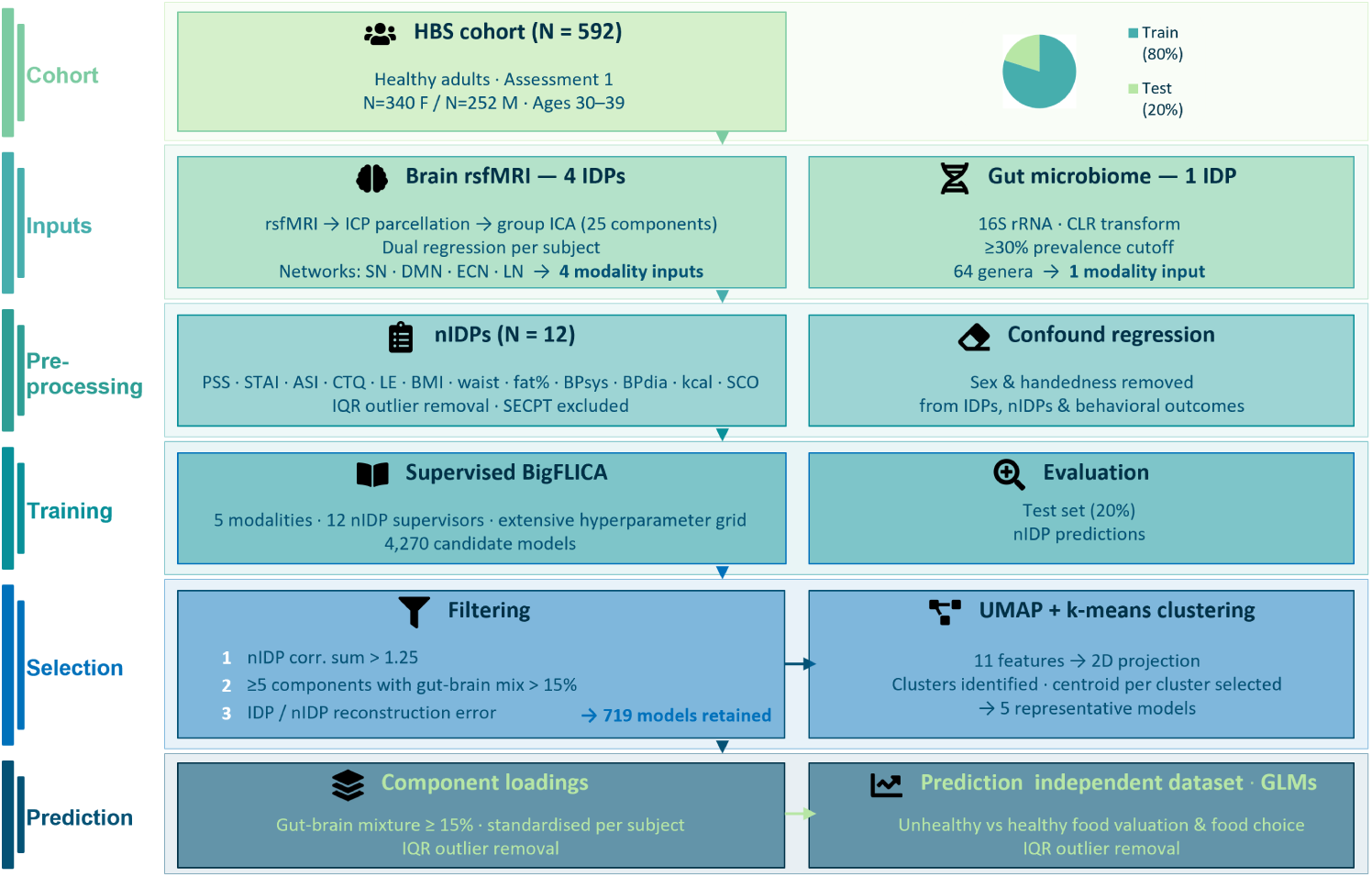
**Overview of the analysis.**

Pre-processing of the modality inputs followed validated pipelines available in https://github.com/silviapapalini/GutBrain-Linkage.

We selected twelve (N = 12) non-imaging-derived phenotypes. Model training was conducted exclusively on the training set using the supervised BigFLICA framework, while the test set was reserved for evaluating generalization performance and model selection. Following initial model training, we implemented a model selection procedure based on predefined quantitative criteria. To identify representative models, we used k-means clustering [54] and plotted the clusters using Uniform Manifold Approximation and Projection (UMAP) [55]. Next, from each cluster, we selected the centroid as representative model for further evaluation. For evaluation, in each selected model, we extracted and standardized subject-specific component loadings from components with meaningful gut–brain mixture (i.e., above 15%) and used them in General Linear Models (GLMs) to predict behavioural performance in an independent FAT (independent measurement within the same participants).

#### 4.3.1 Preparation of IDPs

##### Brain input modalities

Smoothed and denoised Blood-Oxygen-Level Dependent (BOLD) data were obtained directly from the HBS dataset, with full preprocessing details available in the online metadata documentation [20]. Briefly, preprocessing of rsfMRI data followed standard procedures previously applied to the UK Biobank dataset. These included the removal of spatial and temporal artifacts, motion correction, and spatial normalization. All steps were conducted using FSL tools (FMRIB Software Library), as described in the HBS metadata repository.

We first identified functional networks via Instantaneous Connectivity Parcellation (ICP) [56], a method designed to capture fast, transient interactions in brain activity with improved spatial resolution, particularly within subcortical structures. ICP is well-suited to our research focus, as food valuation and decision-making processes involve deep brain areas that are difficult to detect with classical spatial parcellations. These include Insular cortex, ventromedial Prefrontal Cortex, and Orbitofrontal Cortex, all implicated in valuation processes; amygdala, brain-stem, nucleus accumbens (NAc), core nodes of the meso-cortico-limbic system; Anterior Cingulate Cortex, and dorsolateral Prefrontal Cortex, relevant for stress regulation and cognitive control.

Specifically, for each subject, we created a whole-brain grey matter (GM) initialization mask using FSL FIRST on T1-weighted MPRAGE images via the fslanat pipeline [57]. The GM mask used in ICP was thresholded at a Partial Volume Estimate (PVE) = 1, generated using FAST. In the present study, T1 images were aligned to MNI standard space using FLIRT (12 degrees of freedom affine transformation). Next, separately for each hemisphere, we extracted the mean BOLD time series from all voxels within the GM mask using fslmeants. Subsequently, each voxel’s time series was standardized individually.

The standardized mean time series was then multiplied element-wise with each standardized voxel-wise time series, yielding “unfolded” time series that represent moment-to-moment correlations [56]. We then transformed the resulting unfolded time series into MNI space using FSL FLIRT, applying 12 degrees of freedom.

Next, to identify spatially coherent components, the unfolded time series across all subjects were entered into a group-level ICA (25 components) using a temporal concatenation model as implemented in MELODIC [58]. Finally, we used a dual regression analysis to extract individual-level spatial maps and time series for each ICA component [59]. The four brain modalities of interest were represented by expected components of the SN, DMN, ECN, and LN, which were identified via visual inspection (comparing each ICA spatial map to known network templates [59]). These four modalities were then used as inputs for the Linked Independent Component Analysis.

##### Gut microbiome

The gut microbiota data were obtained from the HBS dataset. This included the relative abundance measures obtained through 16S rRNA sequencing. To handle the compositional nature of the data, we applied a centered log-ratio (CLR) transformation. Next, to reduce the zero-inflated nature of the distributions, we retained only bacteria present in more than 70% of the sample (i.e., with fewer than 30% zero values), as in [19]. This resulted in sixty-four (N = 64) bacterial taxa that were included as modality input for the Linked Independent Component Analysis.

In this study, we did not have missing values for the IDPs, since during HBS data request, we requested only data from individuals that had brain and gut variables and at least the presence of heart rate in relation to the SECPT (as the main nIDP, which was, however, not included in the final SuperBigFLICA, see manipulation check of the SECPT).

#### 4.3.2 Preparation of nIDPs

##### Manipulation check of the SECPT

To assess the effectiveness of the stress induction (SECPT), we conducted manipulation checks on three key markers: salivary cortisol, HR, and self-reported subjective stress. These markers were particularly important for the present study given the healthy nature of the sample. For each measure, repeated assessments were collected at three time points (pre-stress, during/after stress, and recovery). Separate one-way repeated-measures ANOVAs were performed with Time (3 levels) as the within-subject factor. Significant main effects were followed up with Tukey’s HSD post hoc tests to examine pairwise time point differences. Participants with missing data at one or more time points were excluded listwise per analysis.

Unexpectedly, the task did not elicit a significant group-level stress response in any of the measures investigated, see supplementary material. Exploring the reasons for this was out of the scope of this project, therefore, differently from what preregistered, we decided to not include these measures (including carotid dilation) into the nIDPs list.

The distributions of the scores from the STAI, PSS, ASI, influence of life events, CTQ, and SCO, as well as those from the kilocalories ingested, body fat percentage, systolic and diastolic blood pressure at rest, BMI, and waist circumference, were inspected and a log transformation was applied only to the scores from the CTQ to reduce its skewness. These variables were included as nIDPs into the model.

#### 4.3.3 Preparation of the behavioural data

The behavioural outcome we aimed to predict is the valuation and choice of unhealthy food vs. healthy items. For the valuation outcome, we analyzed the data from the auction phase of the food task.

During the valuation part of the Food Auction Task, participants viewed single food items and placed a monetary bid (with a maximum of two euros) representing their willingness-to-pay (WTP) for each item. A Becker–DeGroot–Marschak (BDM) style auction procedure was used: a bid was compared to a randomly drawn price, and if the participant’s bid met or exceeded the random price, they won the item and paid that price; otherwise, they kept their endowment and did not receive the item [60, 61]. Following the auction, a food choice part presented participants with pairs of food items (constructed based on individual WTP rankings). On each trial, participants chose between two foods, allowing the measurement of choice behaviour as a function of relative subjective value. This two-stage design enables dissociation of valuation processes (bidding) from choice preferences between alternative food items. We regressed out the effects of sex and handedness from the ratio value between unhealthy vs healthy items from valuation and choice phase. These unconfounded indexes were the two dependent variables we attempted to predict using the subjective loadings from components with gut-brain mixture within classical GLMs.

#### 4.3.4 Model training

The supervised BigFLICA algorithm was used to extract latent components that captured shared variation across multiple brain and microbiota modalities (IDPs), guided by our nIDPs. The model integrated five input modalities: four brain networks, all derived from resting-state fMRI, as well as gut microbiota composition data. Supervision was provided by 12 non-imaging derived phenotypes (nIDPs), including psychological, behavioural, and lifestyle measures. These nIDPs served as guiding variables, steering the decomposition toward components with strong associations with the traits of interest. Components were considered relevant if they showed at least 15% mixture contribution from at least one brain modality and the microbiota modality.

To evaluate model performance, the dataset was randomly divided into a training set (80%) and a test set (20%). The training set was used to fit the supervised BigFLICA model, learning latent components that jointly captured variation across the input modalities while maximizing their predictive association with the nIDPs. The test set was held out during training and used exclusively for model evaluation. For each subject in the test set, we used the trained model to extract subjective component loadings, which were then used to predict the behavioural data obtained from the food auction task. The performance of the model was assessed by computing the correlation between the predicted and actual nIDP values in the test set, providing a measure of the generalizability of the learned components to the unseen data.

Before model training, and as done for the behavioural outcomes, we regressed the effects of gender and handedness from both IDP and nIDP data to remove potential confounding. We then conducted an extensive hyperparameter tuning phase, testing combinations of latent dimensionality (35, 55, 75, or 90 components), learning rates (*α*: 0.01, 0.03, 0.05, or 0.07), reconstruction weights for IDPs (*λ*_1_: 0.55, 0.65, 0.75, or 0.85) and nIDPs (*λ*_2_: 0.1, 0.2, 0.3, or 0.4), and sparsity penalties for IDPs (*λ*_3_: 0.4, 0.6, or 0.75) and nIDPs (*λ*_4_: 0.1, 0.2, 0.3, or 0.4). Beyond this core grid, we extended the search in three directions. First, we probed more extreme latent dimensionalities (20 and 100 components) with learning rates of 0.02, 0.04, and 0.06, excluding combinations with the smallest *λ*_1_ (0.55), which is most critical for reconstructing the gut-microbiota and brain input modalities. Second, we paired an intermediate learning rate (0.035) with high first and third regularization parameters (*λ*_1_ and *λ*_3_ set to 0.85) to capture models with strong reconstruction of both IDPs and nIDPs, using 80, 90, and 110 components across the same *λ*_2_ and *λ*_4_ ranges. Third, we sampled random combinations of comparable parameters to cover some of the remaining model space (see combo parameters.csv in the GutBrain-Linkage GitHub repository). This parameter selection was intended to include a broad range of candidate models while keeping computational and storage costs affordable.

#### 4.3.5 Model evaluation and final model selection

To identify interpretable and biologically relevant models from the hyperparameter tuning phase, we applied a three-stage filtering (arbitrary) procedure based on our specific priorities. First, we excluded models in which the total sum of correlations between the predicted and actual nIDPs in the test set was below 1.25, as these models exhibited very low predictive performance on the supervised traits. Second, we removed models with fewer than 5 components that showed less than 15% of mixture between brain and gut information, ensuring that retained models captured large shared variance across modalities that we could explore. Third, based on the test dataset, we set an arbitrary threshold of 6 on the reconstruction error for the IDPs. This value was selected because the visual inspection of brain reconstruction in models falling above this value was generally poor (i.e., no detectable functional activation). Similarly, a threshold of 7 was applied for the reconstruction error of the nIDPs and the error of their sparseness (test dataset). These thresholds were relatively generous, allowing us to ensure preservation of models that potentially retained a sufficient degree of biological and psychological interpretability. To sum up, this procedure yielded a subset of models (N = 719, surviving count after filtering the total 4270) that performed well across multiple evaluation criteria, including reconstruction fidelity, supervision strength, and multimodal integration.

To further examine this model space and identify representative exemplars, we used UMAP to project the high-dimensional models each characterized by 11 features (number of latent components, learning rates, the four lambda parameters and their respective errors, overall correlation value of all the nIDPs from test-dataset, and the number of components with gut-brain mixture), into a two-dimensional space. We opted for UMAP over Pareto front techniques, since the latter did not yield a clear or visually interpretable solution, likely due to the high dimensionality and complexity of the model performance landscape. We then applied k-means clustering to the 2D UMAP projection to group similar models based on their overall characteristics. For each resulting cluster, we selected the centroid, which served as representative model for the behavioural testing using GLMs.

#### 4.3.6 Statistical analysis for behavioural prediction

From each selected representative model, we extracted subject-specific loadings from the components with meaningful gut–brain mixture and explored whether these (standardized) loadings could predict individual behaviour in the independent food auction task dataset. Specifically, we applied a general linear model to the selected models from each cluster, to explore whether any of these components were associated with both the valuation and choice of unhealthy versus healthy food items (as these two psychological processes are strongly correlated, see the supplementary information in Appendix E). This final evaluation step explored whether latent components reflecting shared gut–brain variance, guided by our nIDPs (e.g., affective-related and body composition traits), could predict high valuation and choice (value-based decision-making) of unhealthy items (our preregistered hypothesis).

#### 4.3.7 Follow-up checks of model robustness

To ensure that the train and test data sets used were statistically comparable in their nIDPs and confounders we ran several *t*-tests to check for significant differences, see the supplementary information in Appendix E. Second, to check the stability of the supervised FLICA model with respect to random initialization, we trained one hundred independent new models (N = 100) using the original train-test split. All other parameters from model number 3280 were kept constant: the number of components was fixed at 100, the learning rate at 0.04, and the maximum number of iterations at 300. Regularization was controlled using the fixed set of lambda values: 0.75, 0.2, 0.6, 0.2. Specifically, from these 100 new models we predicted all the nIDPs and plot the histogram correlations between the predicted and actual nIDP values for both train and test dataset of the nIDPs with higher loadings (ASI, BMI, and life events influence). Third, to check the stability of the supervised FLICA model with respect to random train–test splits, we repeated the same procedure across multiple random data partitions. Model initialization was kept constant across runs, ensuring that variability in outcomes reflected only differences in how the data were partitioned. Fourth, a 5-fold cross-validation was performed to further check the stability and generalizability of the model across different data partitions. The 5-fold cross-validation was performed by partitioning the dataset into five equal folds; in each iteration, four folds were used for training and one for testing. The resulting correlations were averaged across folds and plotted alongside those obtained from the original fold for ASI, BMI, and life events influence, see **Fig. 9c**. Finally, we checked whether component 93 reappeared in similar models included in cluster 3, see table 1.

## Contributions

**Silvia Papalini:** Conceptualization, Data curation, Formal analysis, Investigation, Methodology, Software, Validation, Visualization, Writing original draft, Writing – review & editing. **Esther Aarts:** Conceptualization, Supervision, Writing – review & editing. **Nils Kohn:** Conceptualization, Funding acquisition, Resources, Methodology, Supervision, Writing – review & editing.

## Acknowledgements

The authors thank Michel Doŕe (HBS team member) for his support with data transfer; Jos Seegers (MyMicroZoo) for his input of gut microbiota functioning; Ingredion for helpful feedback on the manuscript.

## Declarations

The authors declare no competing interests.

## Ethics approval and consent to participate

The Healthy Brain Study was approved by the Institutional Review Board of Radboud University Medical Center on 23 May 2019 (reference number: 2018–4894), in accordance with the Declaration of Helsinki. All participants provided written informed consent prior to participation. The current study used data from the HBS cohort under data release number 2024-4; no additional ethical approval was required for secondary analysis of this pre-approved dataset.

## Data availability

The primary data used in this study were obtained from the Healthy Brain Study (HBS) cohort. Researchers can apply for access through the HBS consortium (https://www.healthybrainstudy.nl). Access is subject to data governance procedures of the Radboud University Medical Center. Derived data (standardized subject-specific component loadings from model 3280 used in the GLM analyses) are available from the corresponding author upon reasonable request, subject to applicable data governance constraints.

## Code availability

All analysis scripts are publicly available at: https://github.com/silviapapalini/GutBrain-Linkage.

## Appendix A Additional information

### nIDPs:Manipulation check of the SECPT

#### Cortisol

A significant effect of Time was found for cortisol levels (F(2, 1670) = 6.78, *p*-value *<* 0.001). Post hoc comparisons indicated an unexpected decrease from baseline (Time 1) to post-SECPT (Time 2) (*p*-value = 0.026), followed by a significant increase from Time 2 to Time 3 (*p*-value = 0.001). Unexpectedly, cortisol levels at recovery (Time 3) were not significantly different from baseline (*p*-value = 0.58), indicating return to initial levels.

#### Heart Rate

HR showed a significant Time effect (F(2, 1758) = 5.19, *p*-value = 0.006). Unexpectedly, it decreased significantly from Time 1 to Time 2 (*p*-value = 0.004), with no further significant changes between Time 2 and Time 3 (*p*-value = 0.46) or between Time 1 and Time 3 (*p*-value = 0.11).

#### Subjective Stress

Subjective stress ratings varied significantly over time (F(2, 1700) = 22.94, *p*-value *<* 0.001). Unexpectedly, participants reported lower stress at Time 3 compared to both Time 1 (*p*-value *<* 0.001) and Time 2 (*p*-value *<* 0.001). The change from Time 1 to Time 2 was not statistically significant (*p*-value = 0.26).

Taken together, these results indicate that, in the current subsample, the SECPT did not elicit a consistent or robust stress response across physiological and subjective measures, suggesting an unsuccessful stress induction in this HBS-subsample, see **Fig. A1**.

**Fig. A1:**
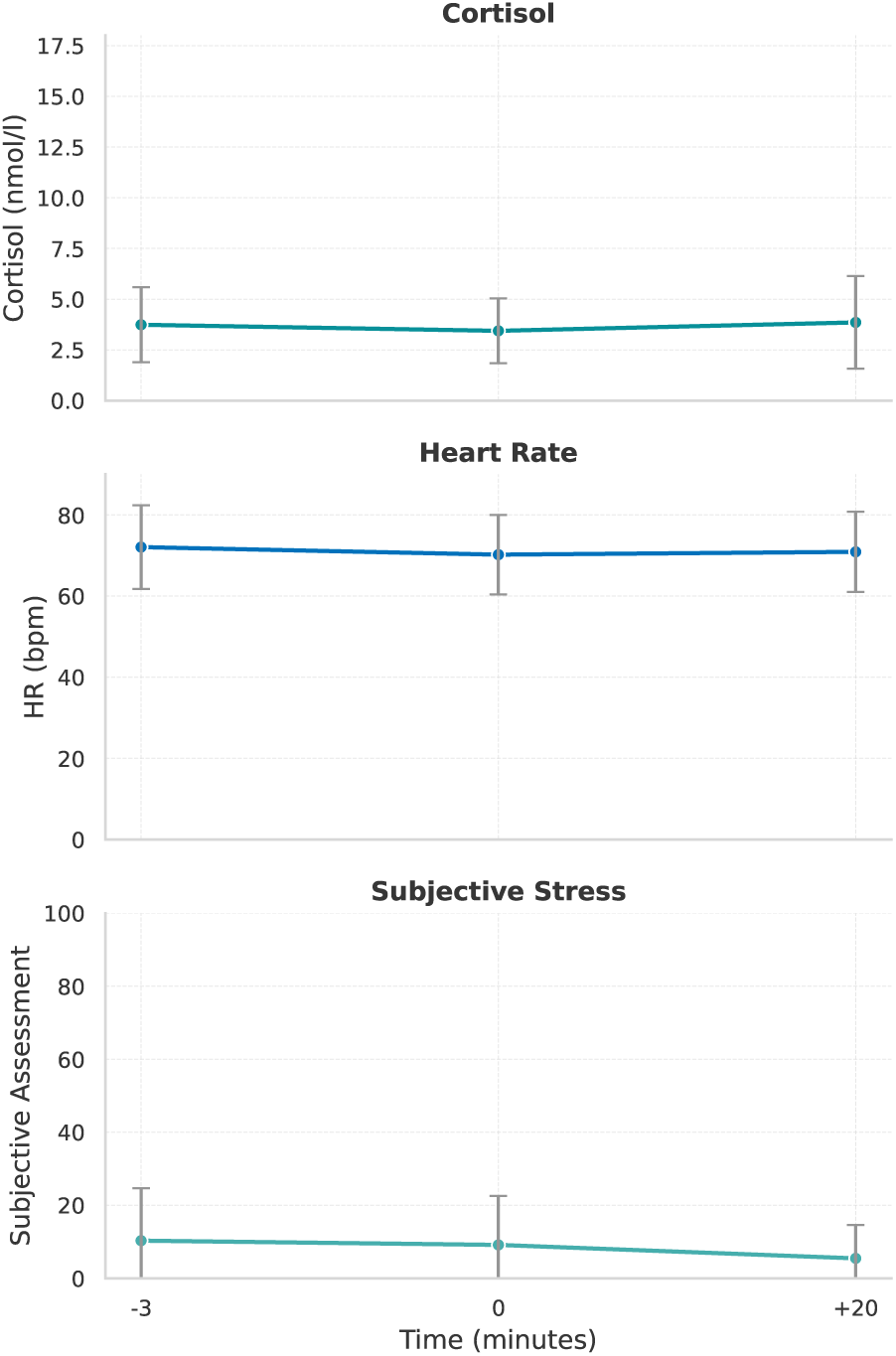
Effect of the SECPT on cortisol levels, heart rate, and subjective stress. Time 0 indicates the conclusion of the task.

## Appendix B Additional information component 93 of model 3280: functional connectivity found in the LN and DMN

**Table B1:** Cluster Peaks from fMRI Analysis: LN.

Appendix B Additional information component 93 of model 3280: functional connectivity found in the LN and DMN.
| Directionality | n Voxels | Cluster | Value | MNI X | MNI Y | MNI Z | Area |
| --- | --- | --- | --- | --- | --- | --- | --- |
| Negative | 25 | 1 | -3.98 | 62.00 | -40.00 | 2.00 | Right Middle Temporal Gyrus |
| Negative | 55 | 2 | -4.05 | 54.00 | -52.00 | 22.00 | Right Angular Gyrus |
| Negative | 50 | 3 | -4.85 | 44.00 | 40.00 | 14.00 | Right Frontal Pole |
| Negative | 23 | 4 | -3.89 | 46.00 | 54.00 | 8.00 | Right Frontal Pole |
| Negative | 26 | 5 | -4.55 | 30.00 | 26.00 | 40.00 | Right Middle Frontal Gyrus |
| Negative | 52 | 6 | -4.65 | 24.00 | -82.00 | -18.00 | Right Occipital Fusiform Gyrus |
| Negative | 58 | 8 | -5.49 | 18.00 | -6.00 | 24.00 | Right Caudate |
| Negative | 34 | 10 | -4.19 | 8.00 | -72.00 | 40.00 | Right Precuneous Cortex |
| Negative | 69 | 11 | -4.38 | 2.00 | -2.00 | 4.00 | Right Thalamus |
| Negative | 71 | 12 | -4.76 | 4.00 | -10.00 | 56.00 | Right Supplementary Motor Cortex |
| Negative | 41 | 13 | -4.41 | -2.00 | -92.00 | 16.00 | Left Occipital Pole |
| Negative | 120 | 14 | -5.68 | -16.00 | 0.00 | 24.00 | Left Caudate |
| Negative | 23 | 15 | -4.47 | -24.00 | -40.00 | -20.00 | Left Temporal Fusiform Cortex |
| Negative | 22 | 16 | -4.93 | -16.00 | -4.00 | -14.00 | Left Amygdala |
| Negative | 27 | 17 | -3.68 | -20.00 | 34.00 | 46.00 | Left Superior Frontal Gyrus |
| Negative | 78 | 18 | -5.20 | -26.00 | -30.00 | -28.00 | Left Parahippocampal Gyrus |
| Negative | 29 | 19 | -5.19 | -30.00 | 46.00 | -10.00 | Left Frontal Pole |
| Negative | 23 | 21 | -4.14 | -32.00 | 8.00 | -16.00 | Left Insular Cortex |
| Negative | 44 | 22 | -5.07 | -54.00 | -12.00 | -10.00 | Left Superior Temporal Gyrus |
| Negative | 20 | 23 | -4.18 | -62.00 | -4.00 | 14.00 | Left Precentral Gyrus |
| Negative | 21 | 24 | -4.02 | -66.00 | -38.00 | -6.00 | Left Middle Temporal Gyrus |
| Positive | 37 | 1 | 4.54 | 46.00 | -68.00 | 0.00 | Right Lateral Occipital Cortex |
| Positive | 38 | 2 | 3.92 | 54.00 | -70.00 | 32.00 | Right Lateral Occipital Cortex |
| Positive | 36 | 5 | 4.56 | 44.00 | 50.00 | -8.00 | Right Frontal Pole |
| Positive | 28 | 7 | 3.88 | 24.00 | 52.00 | 40.00 | Right Frontal Pole |
| Positive | 39 | 8 | 4.56 | 4.00 | -84.00 | 28.00 | Right Cuneal Cortex |
| Positive | 36 | 9 | 4.86 | 2.00 | -66.00 | -2.00 | Right Lingual Gyrus |
| Positive | 22 | 10 | 3.75 | 2.00 | 8.00 | 60.00 | Right Supplementary Motor Cortex |
| Positive | 28 | 11 | 3.92 | 0.00 | -46.00 | 30.00 | Left Cingulate Gyrus |
| Positive | 39 | 12 | 4.68 | -2.00 | 58.00 | 30.00 | Frontal Pole |
| Positive | 22 | 13 | 4.42 | -12.00 | 10.00 | -14.00 | Left Subcallosal Cortex, Putamen, NAc |
| Positive | 46 | 14 | 4.14 | -16.00 | 50.00 | 28.00 | Left Frontal Pole |
| Positive | 26 | 15 | 4.51 | -26.00 | 8.00 | -46.00 | Left Temporal Pole |
| Positive | 20 | 16 | 4.08 | -36.00 | -8.00 | 46.00 | Left Precentral Gyrus |
| Positive | 23 | 17 | 4.15 | -34.00 | -24.00 | 48.00 | Left Precentral Gyrus |
| Positive | 194 | 18 | 4.66 | -52.00 | 8.00 | 34.00 | Left Precentral Gyrus |

**Table B2:**
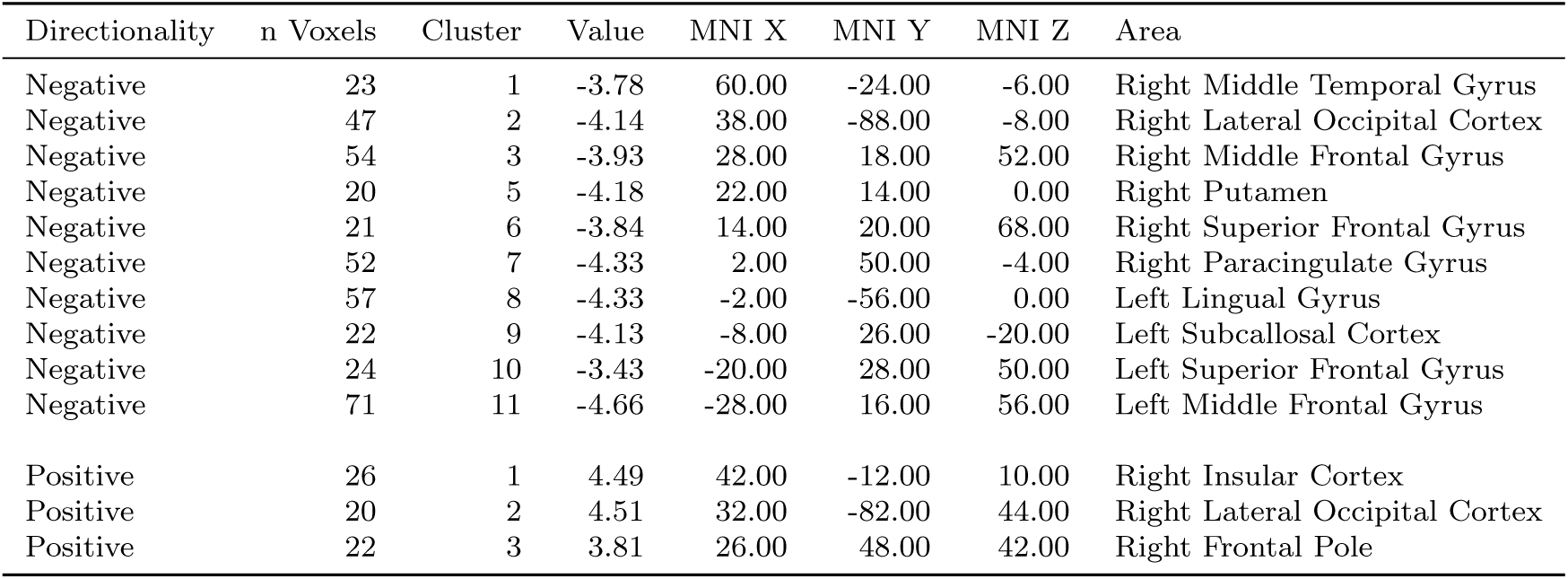
Cluster Peaks from fMRI Analysis: DMN.

## Appendix C Additional information on component 70

Component 70 included positive taxa: Roseburia (*z* = 8.26), Lactobacillus (*z* = 5.58), Lachnospiraceae UCG-008 (*z* = 5.39), Collinsella (*z* = 5.26), Cloacibacillus (*z* = 4.84), Oribacterium (*z* = 4.72), Tyzzerella (*z* = 4.18), Eisenbergiella (*z* = 3.49), Monoglobus (*z* = 3.16), Romboutsia (*z* = 3.03), Fusicatenibacter (*z* = 2.92), Clostridia UCG-014 (*z* = 2.66), Desulfovibrio (*z* = 2.63). And negative taxa: Rikenellaceae RC9 gut group (*z* = *−*2.38), Christensenellaceae (*z* = *−*2.44), CAG-56 (*z* = *−*2.48), Prevotellaceae NK3B31 group (*z* = *−*2.50), Lachnoclostridium (*z* = *−*2.98), vadinBE97 (*z* = *−*2.98), Peptococcus (*z* = *−*3.21), Corynebacterium (*z* = *−*3.48), Sneathia (*z* = *−*3.81), Intestinibacter (*z* = *−*3.91), [Eubacterium] fissicatena group (*z* = *−*4.12), Eggerthella (*z* = *−*4.43), UCG-009 (*z* = *−*4.63), Mogibacterium (*z* = *−*4.78), UCG-002 (*z* = *−*4.97), [Eubacterium] eligens group (*z* = *−*5.13), [Eubacterium] hallii group (*z* = *−*5.48), Slackia (*z* = *−*5.75), Lachnospiraceae NK4A136 group (*z* = *−*6.50), [Eubacterium] siraeum group (*z* = *−*6.80).

## Appendix D Additional information about model 3280: statistical Comparison nIDPS (Train vs. test)

STS: *t* = 0.24, *p*-value = 0.814

PSS: *t* = 1.29, *p*-value = 0.197

kilocal: *t* = 0.10, *p*-value = 0.924

Fat perc: *t* =-0.41, *p*-value = 0.686

Sys BP: *t* = 0.84, *p*-value = 0.403

Dia BP: *t* = 0.87, *p*-value = 0.383

ASI: *t* = 0.27, *p*-value = 0.785

LE Influence: *t* = 0.71, *p*-value = 0.481

BMI: *t* =-0.25, *p*-value = 0.799

Waist Cir: *t* =-0.12, *p*-value = 0.908

SCO: *t* = 1.02, *p*-value = 0.307

CTQlog: *t* = 0.41, *p*-value = 0.682

left/right handed: *t* =-0.30, *p*-value = 0.766

Sex: *t* =-0.14, *p*-value = 0.892

## Appendix E Additional information on FAT

**Table E:**
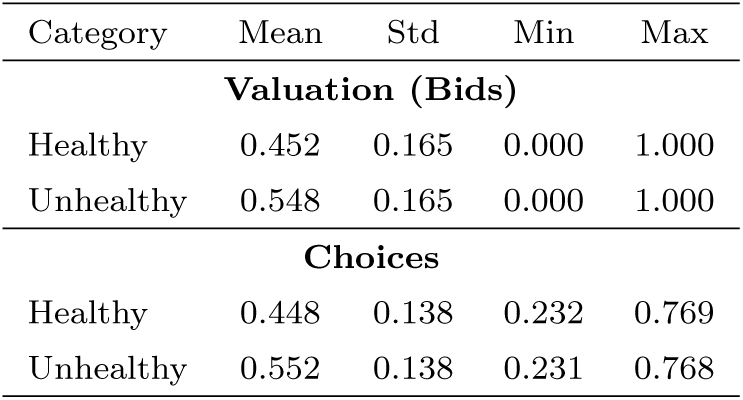
Descriptive statistics for healthy and unhealthy food categories.

| Category | Mean | Std | Min | Max |
| --- | --- | --- | --- | --- |
| **Valuation (Bids)** |  |  |  |  |
| Healthy | 0.452 | 0.165 | 0.000 | 1.000 |
| Unhealthy | 0.548 | 0.165 | 0.000 | 1.000 |
| **Choices** |  |  |  |  |
| Healthy | 0.448 | 0.138 | 0.232 | 0.769 |
| Unhealthy | 0.552 | 0.138 | 0.231 | 0.768 |

**Fig. E1:**
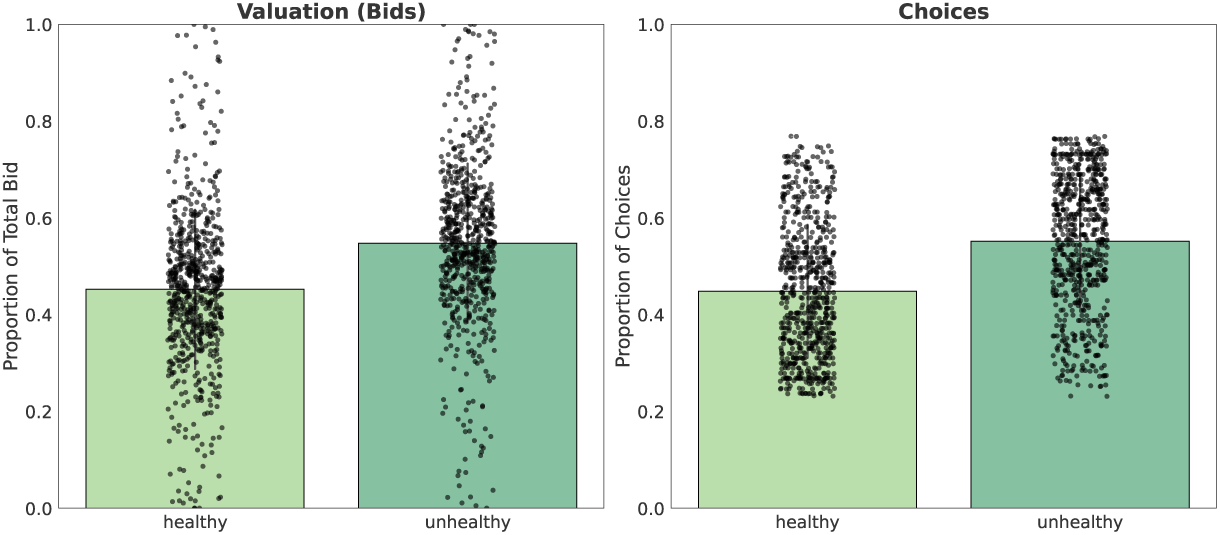
Mean proportion of healthy and unhealthy foods for valuation and choices in the food auction task. Error bars represent standard deviations. Spearman correlation between unhealthy bids and unhealthy choices: *r* = 0.83, *p*-value *<* 0.001. This indicates a strong positive association: participants who valued unhealthy foods more in the auction also chose them more often in the decision-making task.

